# Interpretable deep learning of label-free live cell images uncovers functional hallmarks of highly-metastatic melanoma

**DOI:** 10.1101/2020.05.15.096628

**Authors:** Assaf Zaritsky, Andrew R. Jamieson, Erik S. Welf, Andres Nevarez, Justin Cillay, Ugur Eskiocak, Brandi L. Cantarel, Gaudenz Danuser

**Author notes:** Corresponding author. Assaf Zaritsky, Gaudenz Danuser. Equal contribution (Assaf Zaritsky, Andrew R. Jamieson, Erik S. Welf, Andres Nevarez).

## Abstract

Deep convolutional neural networks have emerged as a powerful technique to identify hidden patterns in complex cell imaging data. However, these machine learning techniques are often criticized as uninterpretable “black-boxes” - lacking the ability to provide meaningful explanations for the cell properties that drive the machine’s prediction. Here, we demonstrate that the latent features extracted from label-free live cell images by an adversarial auto-encoding deep convolutional neural network capture subtle details of cell appearance that allow classification of melanoma cell states, including the metastatic efficiency of seven patient-derived xenograft models that reflect clinical outcome. Although trained exclusively on patient-derived xenograft models, the same classifier also predicted the metastatic efficiency of immortalized melanoma cell lines suggesting that the latent features capture properties that are specifically associated with the metastatic potential of a melanoma cell regardless of its origin. We used the autoencoder to generate “in-silico” cell images that amplified the cellular features driving the classifier of metastatic efficiency. These images unveiled pseudopodial extensions and increased light scattering as functional hallmarks of metastatic cells. We validated this interpretation by analyzing experimental image time-lapse sequences in which melanoma cells spontaneously transitioned between states indicative of low and high metastatic efficiency.

Together, this data is an example of how the application of Artificial Intelligence supports the identification of processes that are essential for the execution of complex integrated cell functions but are too subtle to be identified by a human expert.

## Introduction

Recent machine learning studies have impressively demonstrated that label-free images contain information on the molecular organization within the cell (Christiansen et al., 2018; Ounkomol et al., 2018; Sullivan and Lundberg, 2018; Yuan et al., 2018). These studies relied on generative models that transform label-free to fluorescent images, which can indicate the organization and, in some situations, even the relative densities of molecular structures. Model training was achieved by using pairs of label-free and fluorescence images subject to minimizing the error between the fluorescence ground-truth image and the model-generated image. Other studies used similar concepts to enhance imaging resolution by learning a mapping from low-to-high resolution (Belthangady and Royer, 2019; Fang et al., 2019a; Nehme et al., 2018; Ouyang et al., 2018; Wang et al., 2019; Weigert et al., 2018). Common to all studies is the concept that the complex architecture of a deep convolutional neural network can extract from the label-free or low-resolution cell images unstructured hidden information – also referred to as *latent information* – that is predictive of the molecular organization of a cell or its high-resolution image, yet escapes the human eye.

We wondered whether this paradigm could be applied beyond the prediction of features of cell architecture to the prediction of complex cell states that result from the convergence of numerous structural and molecular signaling factors. We combined unsupervised generative deep neural networks and supervised machine learning to train a classifier that can predict the metastatic efficiency of human melanoma cells.

The power of cell appearance for determining cell functional states has been the basis of decades of histopathology (Beck et al., 2011; Yuan et al., 2012) and it has also been explicitly established in predicting the state of signaling pathways that are directly implicated in the regulation of cell morphogenesis (Bakal et al., 2007; Goodman and Carpenter, 2016; Gordonov et al., 2015; Pascual-Vargas et al., 2017; Scheeder et al., 2018; Sero and Bakal, 2017; Yin et al., 2013). Other studies used deep neural networks to classify cell cycle states and diabetic retinopathy from fluorescent-labeled cells (Eulenberg et al., 2017), to predict a differentiation marker prior to the actual expression in the cells from live bright-field microscopy (Buggenthin et al., 2017; Orth et al., 2017), and to reconstruct pseudo-lineages from single cell snapshots (Yang et al., 2020).

Whether morphological cues are also informative of the broader spectrum of cell signaling programs that drive hallmark functions in metastatic cells such as shifts in the regulation of metabolism, cell cycle progression, or cell death is less clear, although very recent work, using conventional shape-based machine learning of fluorescently labeled cell lines, suggests this may be the case (Wu et al., 2020).

The paradigm of extracting latent information via deep convolutional neural networks from label-free and time-resolved image sequences holds particularly strong promise for a task of this complexity, as the design of metrics of cell appearance that encode the state of, e.g., a pro-survival signal, exceeds human intuition. The flip side of learning non-intuitive features is the discomfort of relying on the classification by a ‘black box’ algorithm with poorly interpretable behavior. Especially in a clinical setting, the lack of a straightforward meaning of the classifier determinants is a widely perceived weakness of deep learning systems. By generating “in silico” cell images that were never observed experimentally and by exploiting temporal information from live cell imaging experiments we “reverse engineered” the physical properties of the latent image information that discriminates melanoma cells with low versus high metastatic efficiency. These results not only demonstrate that the internal encoding of latent variables in a deep convolutional neural network can be mapped to physical entities predictive of complex cell states, but they highlight more broadly the potential of “interpreted artificial intelligence” in augmenting investigator-driven analysis of cell functions with an entirely novel set of hypotheses.

## Results

### An assay for live imaging of cell lines and patient-derived xenograft (PDX) melanoma cells

To test whether the latent information extracted from label-free live cell movies can predict the metastatic propensity of melanoma, we relied on a previously established patient-derived xenotransplantation (PDX) assay, in which stage III melanoma were originally extracted from lymph node metastases and repeatedly transplanted between immuno-compromised mice (Quintana et al., 2012). All tumors grew and eventually developed metastases again. However, whereas some tumors showed widespread metastases in various distant organs, referred to as a PDX with high metastatic efficiency, other tumors exhibited only lung metastases, referred to as a PDX with low metastatic efficiency. Even after a decade of transplantation, the PDX models maintain a high level of correlation to metastatic outcome in the human patient. Low efficiency PDXs originated from patients that were cured after surgery and chemotherapeutic treatment.

High efficiency PDXs originated from patients with fatal outcome (Quintana et al., 2012). For this study, we had access to a panel of 9 PDXs, seven of which had known metastatic efficiency and matching patient outcome. For the remaining 2 PDX the metastatic efficiency, including patient outcome, was unknown (Table S1). To define the genomic states of the PDXs with known metastatic efficiency, we sequenced a panel of ∼1400 clinically actionable genes and found them to span the genomic landscape of melanoma mutations, including mutations in BRAF (5/6), CKIT (2/6), NRAS (1/6), TP53 (2/6), and copy number variation (CNV) in CDKN2A (6/6) and PTEN (3/6) (Hayward et al., 2017; Hodis et al., 2012) (Table S2). For one PDX (m528), we were unable to generate sufficient genomic material for sequencing, although the cell culture was sufficiently robust for the single cell imaging assay.

In order to prevent morphological homogenization and to better mimic the collagenous ECM of the dermal stroma, we imaged cells on top of a thick slab of collagen. The cells were plated sparsely to focus on cell-autonomous behaviors with minimal interference from interactions with other cells (Methods). For each plate, we recorded with a 20X/0.8NA lens phase contrast movies of at least 2 hours duration, sampled at 1 minute intervals (Fig. 1A, Video S2). Each recording sampled 10-20 randomly distributed fields of view from 1-4 plates of different cell types, each containing 8-20 individual cells.

**Figure 1:**
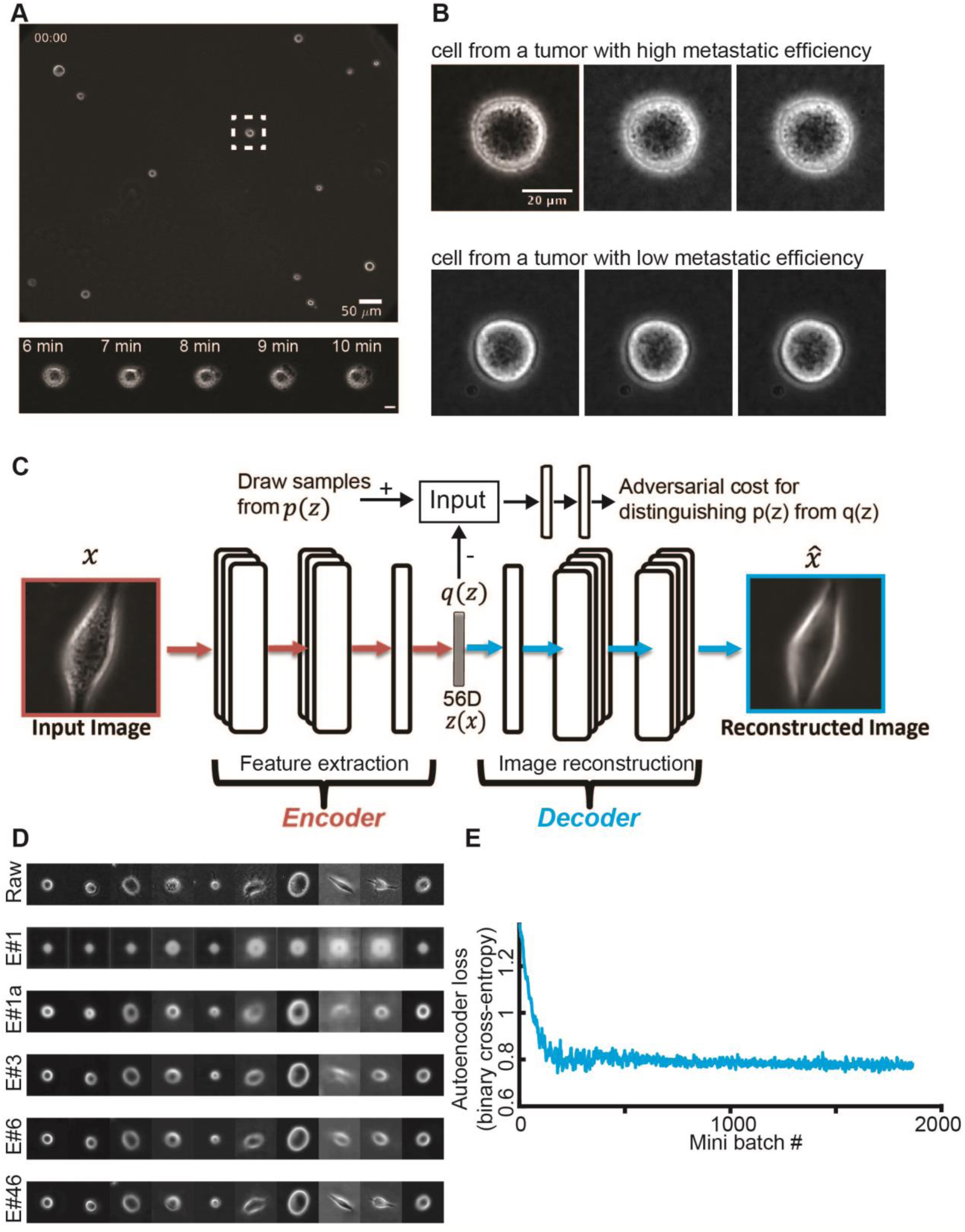
Unsupervised learning of a latent vector that encodes characteristic features of individual melanoma cells. (**A**) Top: Snapshot of a representative field of view of m481 PDX cells. Scale bar = 50 μm. Bottom: Time-lapse sequence of a single cell undergoing dynamic blebbing. Scale bar = 50 μm. (**B**) Representative time-lapse images of single cells from PDX tumors exhibiting low (m498) and high (m634) metastatic efficiency. Sequential images were each acquired 1 minute apart. (**C**) Design of the adversarial autoencoder, comprising an encoder (dark red) to extract from single cell images a 56-dimensional latent vector, so that a decoder can reconstruct from the vector a similar image. The “adversarial” component (top) penalizes randomly generated latent cell descriptors q(z) that the network fails to distinguish from latent cell descriptors drawn from the distribution of observed cells p(z). (**D**) Examples of cell reconstructions. Raw cell images (top): epoch #1 (trained on 10,000 images), epoch #1a (after 1,000,000 images), epoch #3, epoch #6, and epoch #46. (**E**) Convergence of autoencoder loss (binary cross-entropy). Epoch is a full data set training cycle that consists of ∼1.7 million images. Mini-batch is the number of images processed on the GPU at a time. Each mini-batch includes 50 cell images randomly selected for each network parameter learning update. For every epoch, the images order is scrambled and then partitioned into ordered sets of 50 for each mini-batch.

We complemented the PDX data set with equivalently acquired time-lapse sequences of 2 untransformed melanocyte cell lines and 6 melanoma cell lines. The former served as a control to test whether the latent information allows at minimum the distinction of untransformed and metastatic cells. The latter served as a control to test whether the latent information allows the distinction of different cell populations, which, by the long-term selection of passaging in the lab, likely have drifted to a distinct spectrum of functional states than PDX cell populations exhibit.

In total, our combined data set comprises time-lapse image sequences of more than 12,000 single melanoma cells, resulting in approximately 1,700,000 raw images. The cells were typically not migratory but displayed variable morphology and local dynamics (Video S1). Many of the cells were characterized by an overall round cell shape and dynamic surface blebbing (Fig. S1, Fig. S2, Video S2), regardless of whether they belonged to the melanoma group with high or low metastatic efficiency (Fig. 1B qualitatively, quantitative results shown in later figures), which is consistent with reports of primary melanoma behavior *in vivo* (Pinner and Sahai, 2008; Sadok et al., 2015; Sahai and Marshall, 2003) and on soft substrates *in vitro* (Cantelli et al., 2015; Welf et al., 2016). Thus, we speculated that cell shape or motion might not be informative of the functional state of a melanoma cell. Nonetheless, we still noted significant textural variation and dynamics within the phase contrast images. Thus, we wondered whether these images contain visual unstructured cell appearances that could predict the functional cell state.

### Design of adversarial autoencoders for unsupervised feature extraction

After detection and tracking of single cells over time (Methods), we used the cropped single cell images as atomic units to train an adversarial autoencoder (Makhzani et al., 2015) (Fig. 1C, Methods). The autoencoder comprises a deep convolutional neural network to “encode” the image data of a single cell in a vector of latent information, from which a structurally symmetric deep convolutional neural network “decodes” synthetic images (Fig. 1C). The networks are trained to minimize the discrepancy between input and reconstructed images. The adversarial component penalizes randomly generated latent cell descriptors q(z) that the network fails to distinguish from latent cell descriptors drawn from the distribution of observed cells p(z), thus ensuring regularization of the latent information space. Our network architecture employed the part of the network originally used to reconstruct landmarks of the cell nucleus and cytoplasm in (Johnson et al., 2017). Although we supplied the network with phase-contrast melanoma cell images instead of fluorescence images, the adversarial autoencoder displayed fast convergence in reconstructing phase-contrast like cell images (Fig. 1D-E, Video S3, Fig. S3). Importantly, the network training is entirely agnostic to the subsequent classification task. The goal of this step was merely to determine for each melanoma cell an unsupervised latent cell descriptor that holds a compressed representation of a cell’s input image for further classification of melanoma cell states.

### The adversarial autoencoder latent vector is a *quantitative* measure for cell appearance

We verified that the 56-dimensional latent vector defines a quantitative measure for cell appearance, i.e., increasing distances between two data points in the latent space correspond to increasing differences between the input images. We first validated that variations in the latent vector cause variations in cell appearances (Fig. 2A). To accomplish this we numerically perturbed the latent vector after encoding a cell image with varying amounts of noise and calculated the mean squared error between the raw and reconstructed images. As expected, the mean squared error between reconstructed and raw images monotonically increased with increasing amount of noise added in the latent space (Fig. 2B). Hence, the trained encoder generates a locally differentiable latent space. Second, we interpolated a linear trajectory in the latent space between two experimentally observed cells, as well as between two random points, and confirmed, visually and quantitatively, that the decoded images gradually transform from one image to the other (Fig. 2C-D, Video S2Morphing, Fig. S4). Hence, the trained encoder generates a latent space without discontinuities. Third, we calculated the latent space distances between a cell at time *t* and the same cell at *t*+100 minutes and between a cell at time *t* and a neighboring cell in the same sample at time *t*. The distances between time-shifted latent space vectors for the same cell were significantly shorter than those between neighboring cells (Fig. 2E). Hence, the combined effects of time variation in global imaging parameters and of morphological changes on displacements in the latent space tend to be smaller than the difference between cells, confirming -- like the two previous tests -- that the trained adversarial autoencoder latent space defines a faithful metric for the comparison of melanoma cell appearance in phase-contrast images.

**Figure 2:**
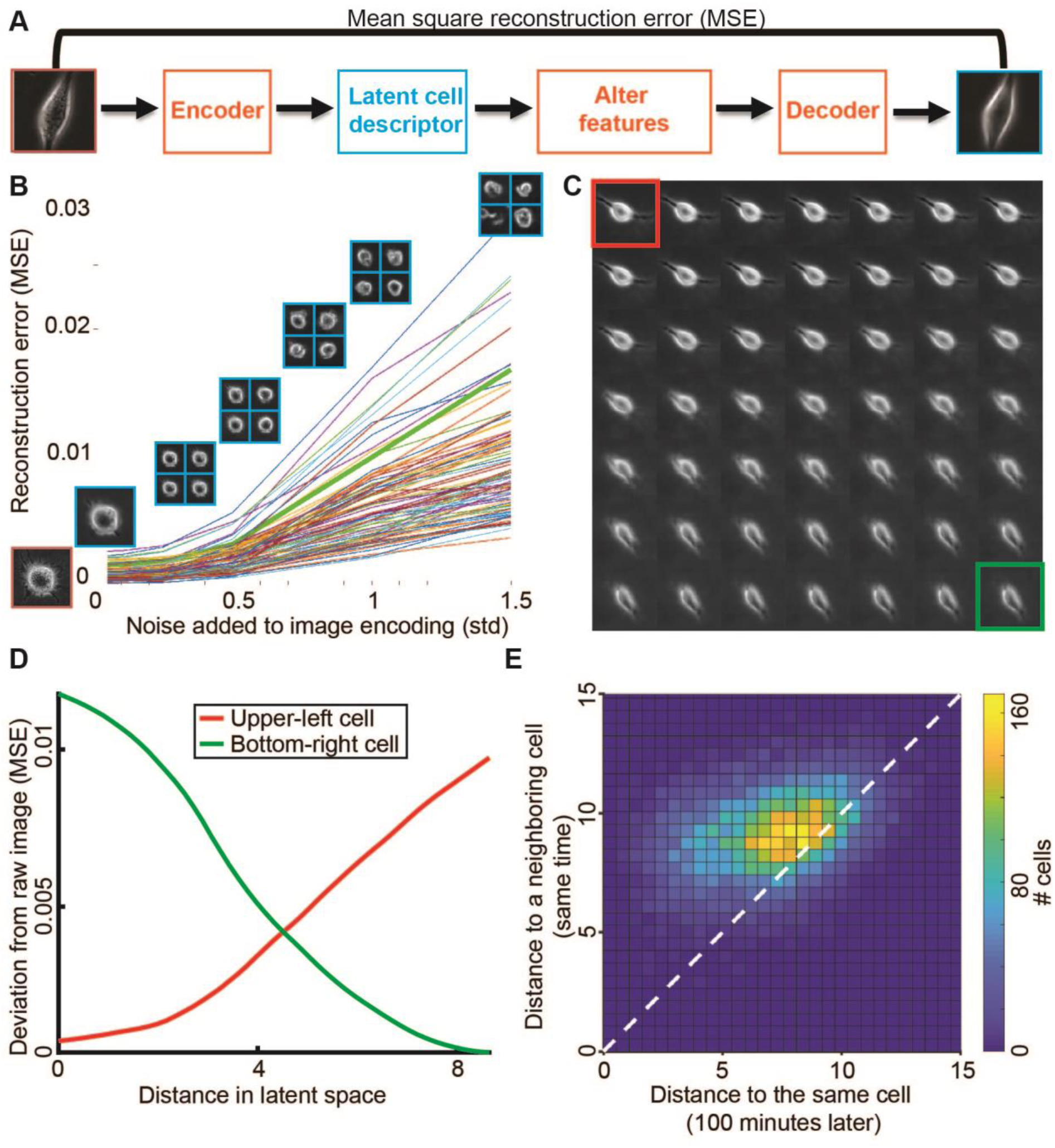
Validation of adversarial autoencoder latent space as a quantitative measure of cell appearance. (**A**) Pipeline to test that increasing shifts in the latent vector of a cell relate to a monotonically increasing shift in cell appearances. (**B**) Increasing perturbation of a particular cell’s latent space vector by Gaussian noise yields an increased deviation of the reconstructed cell image from the original image (image indicated at x = 0). For each noise level, except level 0, four representative reconstructed images are shown. Lines indicate the reconstruction error for 92 randomly selected cells from different cell types and different biological replicates. (**C**) Cell “morphing”. Latent space interpolation shows that a gradual linear transition in latent space yields gradual transition in image space. By “gradual linear transition in latent space” we refer to constant size shifts in feature space for each shift. The trajectory goes from top-left (red) to bottom-right (green). (**D**) Differences of images in panel C and to the start-(red) and endpoint (green) images. (**E**) Cells are more self-similar over time than two neighboring cells at the same time. Two-dimensional histogram of the Euclidean distance between the latent space descriptors of a cell at time 0 and time 100 (x-axis) versus the distance of the same cell to its closest neighboring cell in the same field of view, also at time 0 (y-axis).

### Batch effects (inter-day variability) mask the functional cell state

Batch effects are a major hurdle in the classification of data sets that are acquired over multiple experimental repeats (Boutros et al., 2015; Caicedo et al., 2017). In the case of the presented label-free imaging assay, such effects may arise from uncontrolled experimental variables such as variations in the properties of the collagen gel, illumination artifacts, or inconsistencies in the phase ring alignment between sessions. Autoencoders are known to be very effective in capturing subtle image patterns. Therefore, they may pick up batch effects that mask image appearances related to the functional state of a cell. For a data set free of batch effects, and under the assumption that intra-patient/cell line variability in image appearance is less than inter-patient/cell line appearance, we expect the latent cell descriptors of the same cell class on different days to be more similar than the descriptors of different cell classes imaged on the same day.

To test how strong batch effects may be in our data, we simultaneously imaged four different PDXs in an imaging session that we replicated on different days. Every cell was represented by the time-averaged latent space vector over the entire movie. We then computed the Euclidean distance as a measure of dissimilarity between descriptors from the *same* PDX imaged on *different days* to the distribution of Euclidean distances between *different* PDXs imaged on the *same* day (Fig. 3A). For three of the four tested PDXs we could not find a clear difference between the intra-PDX/inter-day similarity and the intra-day/inter-PDX similarity (Fig. 3B).

**Figure 3:**
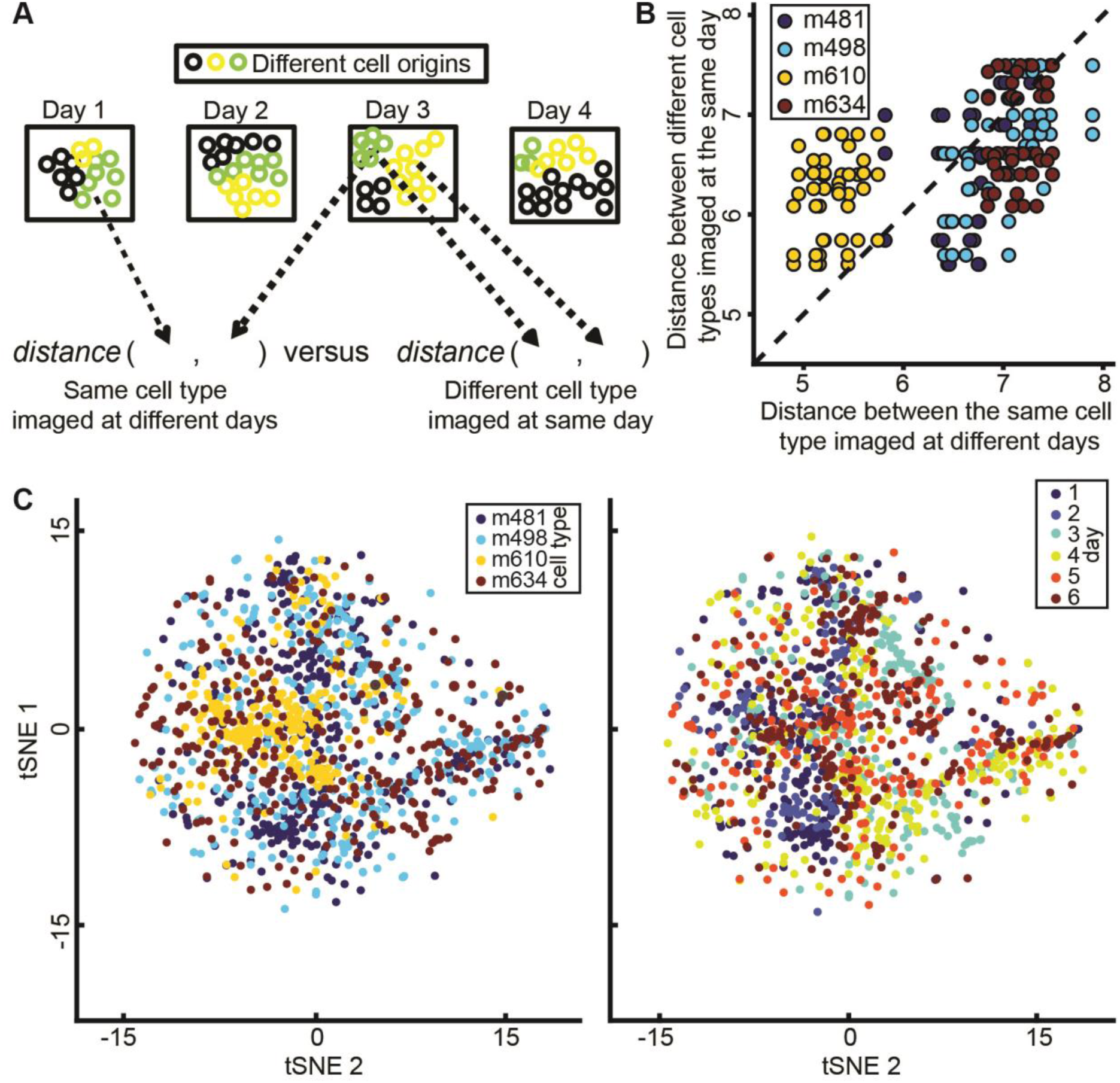
Determining batch effects (day-to-day variability). Cells from four melanoma PDXs (m481, m498, m610, m634) were imaged in one batch, and this experiment was repeated on 6 different days. (**A**) Assessing the distance among different days for the same PDX versus the distance among the different PDXs imaged on the same day. (**B**) Intra-PDX/inter-day distance (x-axis) versus intra-day/inter-PDX distance (y-axis). Each dot represents the distance between the mean time-averaged latent cell descriptors averaged over all cells, arbitrary units. (**C**) tSNE projection of latent space cell descriptors of different PDXs on the same day (left) and of one PDX imaged on different days (right).

Only PDX m610 displayed greater intra-PDX/inter-day similarity than intra-day/inter-PDX similarity. Consistent with this assessment, visualization of all time-averaged cell descriptors over all PDXs and days using PCA (Jolliffe, 2011) or tSNE (Maaten and Hinton, 2008) projections neither showed cell clusters associated with different PDXs nor with different imaging days, except for m610 (Fig. 3C, Fig. S5). These results suggest that the latent space cell descriptors are significantly distorted by batch effects or lack of information on distinct functional states between PDXs.

### The latent cell descriptor can discriminate between different cell types

To overcome the putative batch effects, we sought to transform the auto-encoder latent space into a classifier space that is invariant to inter-day confounding factors, but discriminates between different cell types. This was accomplished by training supervised machine learning models using Linear Discriminant Analysis (LDA). We validated the models in multiple rounds of training and testing, each round with the imaging data of one cell type designated as the test-set, while the rest of the data was used as the training set (Fig. 4A). Hence, the discriminative model is trained with information completely independent of the cell type it is tested on (Jones, 2019). The number of cells from each label was balanced during training to eliminate sampling bias. To overcome the limited statistical power due to the small number of cell types, we also considered the combination of data from one cell type imaged in one day as the test dataset (a single observation). In this case, the training dataset included the remainder of all imaging data, except cells imaged on the same day or the same cell type (Fig. S6).

**Figure 4:**
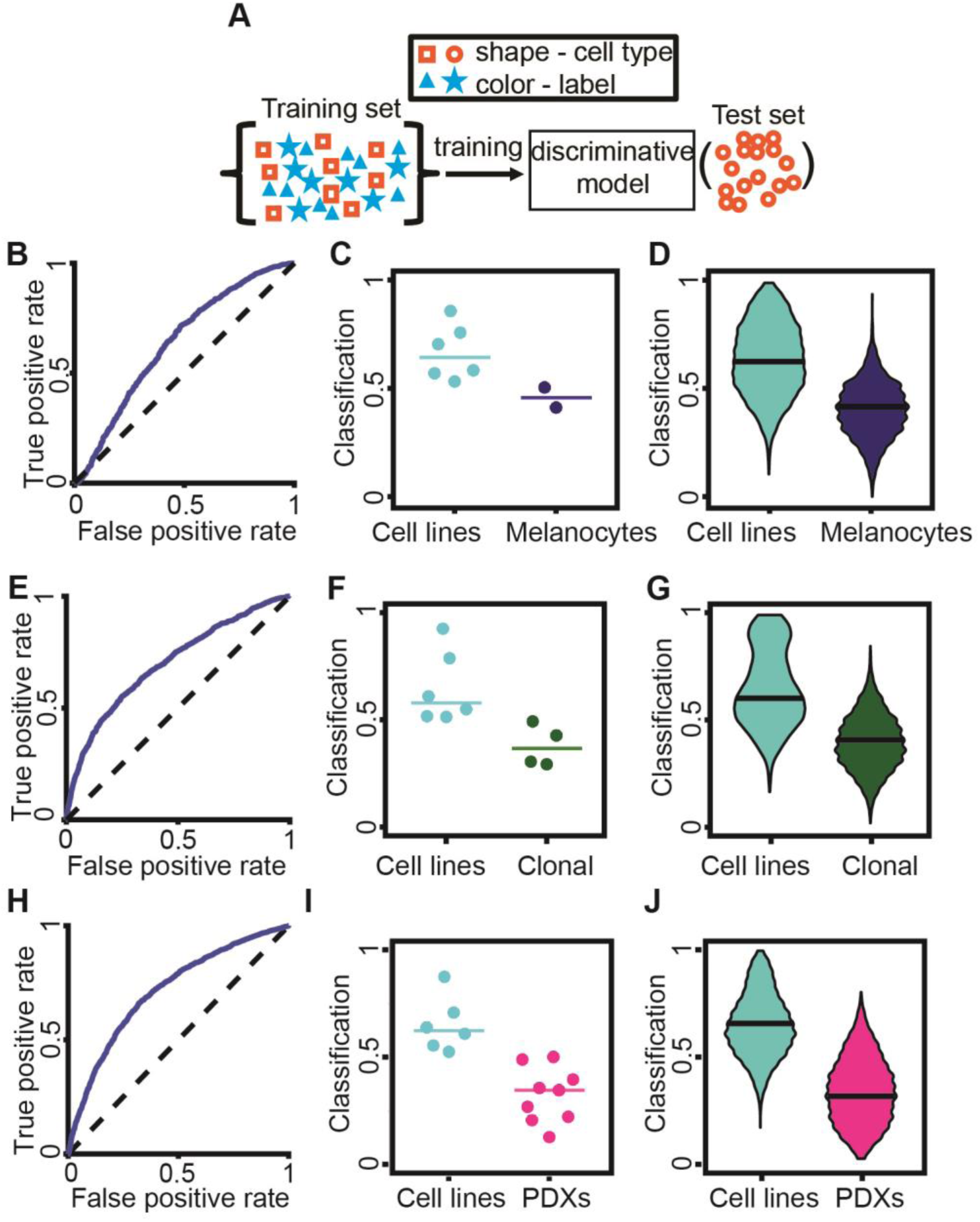
Discrimination of different melanoma cell types. (**A**) Blinding the cell type. Multiple rounds of training and testing were performed. In each round, data from one cell type was used as the test dataset, defining a single observation that was composed of many single cell classifications. The training set contained the rest of the data relevant for the task (e.g., all melanoma cell lines and all PDX when discriminating these two classes). The trained model was completely blind to the cell type used in each test set. The trained model classified each cell in the test set. (**B**) Receiver-Operator Characteristic (ROC) curve for the distinction of the label [melanoma] ‘cell lines’ from the label ‘melanocytes’ [line]. AUC = 0.635. (**C**) Accuracy in predicting the label ‘cell lines’ for a single cell as opposed to the label ‘melanocytes’. Each data point indicates the outcome (fraction of cells classified as ‘cell line’) of testing the cells of one melanoma cell line or melanocyte line. N = 8: 6 melanoma cell lines, 2 melanocyte lines. 7/8 successful predictions. Wilcoxon rank-sum and Binomial statistical tests on the null hypothesis that the classifier scores of a cell line and of melanocytes are drawn from the same distribution, p = 0.071 (Wilcoxon), p = 0.035 (Binomial). (**D**) Bootstrap distribution for predicting the ‘cell lines’ label. For each cell type, we generated 1000 observations by repeatedly selecting 20 random cells and recorded the fraction of these cells that were classified as ‘cell lines’. Horizontal line – median. Wilcoxon rank-sum test p < 0.0001. This analysis demonstrated the ability to discriminate cell lines versus melanocytes from samples of 20 random cells. (**E**) ROC curve for the distinction of the label [melanoma] ‘cell lines’ from the label ‘clonal’ [expansion line]. (**F**) Accuracy in predicting the label ‘cell lines’ for a single cell as opposed to the label ‘clonal’. Each data point indicates the outcome of testing the cells of one melanoma cell line or clonal expansion line. N = 10: 6 melanoma cell lines, 4 clonal expansion lines. 10/10 successful predictions. Wilcoxon rank-sum and Binomial statistical test on the null hypothesis that classifier scores of a cell line are distinct from those of melanocytes p = 0.010 (Wilcoxon), p < 0.001 (Binomial). (**G**) Bootstrap distribution for predicting the ‘cell lines’ label. See panel D. Horizontal line - median. Wilcoxon rank-sum test p < 0.0001. (**H**) ROC curve for the distinction of the label [melanoma] ‘cell lines’ versus the label ‘PDXs’. AUC = 0.714. (**I**) Accuracy in predicting the label ‘cell lines’ for a single cell as opposed to the label ‘PDXs’. Each data point indicates the outcome of testing the cells of one melanoma cell line or PDX. N = 15: 6 cells lines, 9 PDXs. 14/15 successful predicted observations. Wilcoxon rank-sum and Binomial statistical test on the null hypothesis that classifier scores of cell lines and of PDXs are drawn from the same distribution, p < 0.0004 (Wilcoxon), p < 0.0005 (Binomial). (**J**) Bootstrap distribution for predicting the ‘cell lines’ label. See panel D. Horizontal line – median. Wilcoxon rank-sum test p < 0.0001. For all panels we used the time-averaged latent space vector over the entire movie as a cell’s descriptor.

This approach was successful in discriminating transformed melanoma cell lines from non-transformed melanocyte cell lines (Fig. 4B-D, Fig. S7), melanoma cell lines from clonal expansions of these cell lines (Fig. 4E-G, Fig. S8, Methods), and melanoma cell lines from patient-derived xenografts (PDX) (Fig. 4H-J, Fig. S9). We also demonstrated that most pairs of different cell types could be discriminated from one another (Fig. S10). Altogether, these results established that the latent cell descriptor captures information on the functional cell state that is distinct for different cell types.

### Incorporating temporal information to distinguish between cell lines and PDX tumors

To compare the performance of the deep-learned cell descriptors to conventional, shape-based descriptors of cell states (Bakal et al., 2007; Goodman and Carpenter, 2016; Gordonov et al., 2015; Pascual-Vargas et al., 2017; Scheeder et al., 2018; Sero and Bakal, 2017; Yin et al., 2013) we segmented phase contrast cell images of multiple cell types with diverse appearances. We used LEVER (Winter et al., 2016) for this task (Fig. S11) and extracted 13 basic shape features for each cell (Methods). At the same time, we also wondered whether the discrimination of melanoma cell lines from PDXs would benefit from explicit incorporation of temporal information. To do so, we compared three types of simple descriptors derived from shape-based and autoencoder latent space-based cell features, respectively (Fig. S12A). The first relied on cell appearance in single time points, ignoring time information. The second employed the cell trajectories for averaging, which cancels noise for cells with a stationary appearance. The third accounted for switches in cell appearance as the distribution of discrete states a cell visits over time (BOW - “bag-of-words” representation (Sivic and Zisserman, 2009)) (Methods).

We found that purely shape-based descriptors could not distinguish cell lines from PDXs (Fig. S12B). This indicates that the autoencoder latent space captures information from the phase-contrast images that is missed by the shape features. Incorporation of temporal information, especially the time-averaging, slightly (but significantly) boosted the classification performance of LDA models derived from latent space cell descriptors (Fig. S12C). This outcome is consistent with computer vision studies concluding that explicit modeling of time may lead to only marginal gains in classification performance. Based on these findings we used the time-averaged latent space cell descriptors as the basic feature set for cell classification throughout the remainder of our study.

### Live cell histology for classification of melanoma metastatic efficiency

Equipped with the latent space cell descriptors and LDA classifiers, we tested our ability to predict the metastatic efficiency of xenotransplanted melanoma stage III PDXs. Standard cell biology assays, such as measuring the extracellular acidification rate, oxygen consumption rate, and proliferation rate could not discriminate between low and high metastatic efficient PDXs (Fig. S13). In contrast, our image-based approach was able to perfectly discriminate between high- and low-metastatic efficient tumors (Fig. 5B-D). We were also successful at distinguishing PDXs with low versus high metastatic efficiency that were imaged on a single day (small n), by classifiers that were blind to the PDX and to the day of imaging (Fig. S6, Fig. 5E-G, Fig. S14A-B). Cell shape information (Fig. S14C) and mean square displacement analysis of trajectories (Fig. S14D) could not stratify these PDXs. Classifiers trained with the latent space cell descriptor were robust to artificial blurring (Fig. 5H), and illumination changes (Fig. 5I). These results established the potential of the proposed imaging and analytical pipeline as a diagnostic approach, referred to as Live Cell Histology.

**Figure 5:**
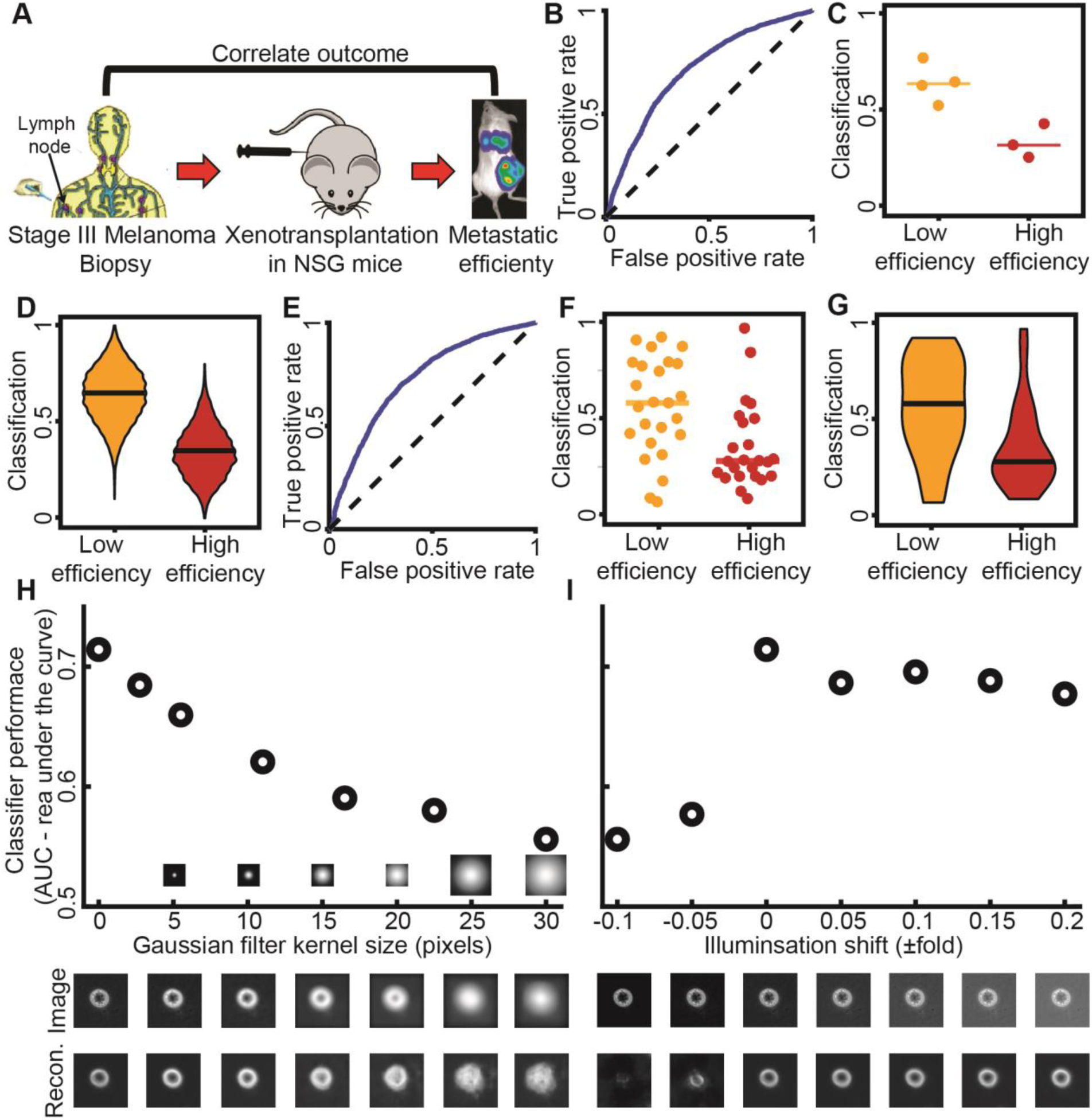
Live Cell Histology can discriminate between PDXs with low versus high metastatic efficiency. (**A**) Correlating outcome in patients and mice (Quintana et al., 2012). (**B**) Receiver Operating Characteristic (ROC) curve. AUC = 0.71. (**C**) Accuracy in predicting for a single cell the label ‘low efficiency’ as opposed to the label ‘high efficiency’ (i.e., the fraction of cells classified as ‘low’). Each data point indicates the outcome of testing the cells of one PDX. N = 7: 4 low efficiency, 3 high efficiency metastasizers. 7/7 predictions are correct. Wilcoxon rank-sum test p = 0.0571. Binomial statistical test p ≤ 0.00782. (**D**) Bootstrap distribution for predicting the ‘low efficiency’ label. For each PDX we generated 1000 observations by repeatedly selecting 20 random cells and recorded the fraction of these cells that were classified as ‘low efficiency’. Horizontal line - median. Wilcoxon rank-sum test p < 0.0001. This analysis demonstrated the ability to predict metastatic efficiency from samples of 20 random cells. (**E-G**) Discrimination results using classifiers that were blind to the cell type and day of imaging (Fig. S6, more observations, smaller n - number of cells for each observation). (**E**) Receiver Operating Characteristic (ROC) curve; AUC = 0.723. (**F**) Accuracy in predicting the label ‘low efficiency’ for a single cell as opposed to the label ‘high efficiency’. Each data point indicates the outcome of testing the cells of one PDX on a particular day. N = 49: 25 low metastatic efficiency, 24 high metastatic efficiency. 32/49 predictions were correct. Wilcoxon rank-sum test p = 0.0042. Binomial statistical test p ≤ 0.0222. (**G**) Bootstrap distribution for predicting the ‘low efficiency’ label. See panel D. Horizontal line - median. Wilcoxon rank-sum test p < 0.0001. (**H**) Robustness of classifier against image blur. Blur was simulated by filtering the raw images with Gaussian kernels of increased size. The PDX m528 was used to compute AUC changes as a function of blur. Representative blurred image (middle) and its reconstruction (bottom). (**I**) Robustness of classifier to illumination changes. AUC as a function of altered illumination (top). Representative image of m528 cell after simulated illumination alteration (middle), and its reconstruction (bottom).

While time-averaged latent single cell descriptors were sufficient to discriminate high and low efficiency metastasizers (Fig. 5), we wondered whether cell “plasticity”, i.e., the capability of a cell to switch between states, would offer additional information on the metastatic state (Pandya et al., 2017). To test this possibility we followed the classifier prediction of an individual cell over time (Methods). For each PDX, we calculated the average rate for a cell to transition between the predicted low and high efficiency state and the fraction of cells that undergo such transitions. Assuming that cell plasticity and predicted metastatic efficiency are unrelated, one would expect that PDXs with a clear-cut prediction of high or low efficiency will be deemed less plastic than PDXs with classifier scores that are close to the decision line of 0.5. Unexpectedly, we found a substantial correlation between both the transition rate as well as the fraction of “plastic” cells and the time-averaged classifier score (Fig. S15). This suggests that highly metastatic PDXs display more cell movement in the latent space, i.e. more variation in the cell appearances over time. The one exception to this rule was PDX m610, which had the lowest classification accuracy. Accordingly, many of its cells located close to the classifier’s decision line, causing random transitions between the low and high efficiency metastatic states. Our data corroborate the notion that the plasticity is a regulated rather than a random behavior.

### Identification of classification-driving features in autoencoder latent space

Our results thus far established the predictive power of the GAN-based, deep-learned latent cell descriptor for the diagnosis of metastatic potential. However, the power of these deep networks to recognize statistically meaningful image patterns that escape the attention of a human observer is also its biggest weakness (Belthangady and Royer, 2019; Caicedo et al., 2017): What is the information extracted in the latent space that drives the accurate classification of low versus high metastatic PDXs? When we plotted a series of cell snapshots from one PDX in rank order of the classifier score, there was no pattern that could intuitively explain the score shift (Fig. 6A). This outcome was not too surprising given that much of the cell appearance is likely unrelated to metastasis-enabling functions, including the image signals associated with batch effects (Fig. 3).

**Figure 6:**
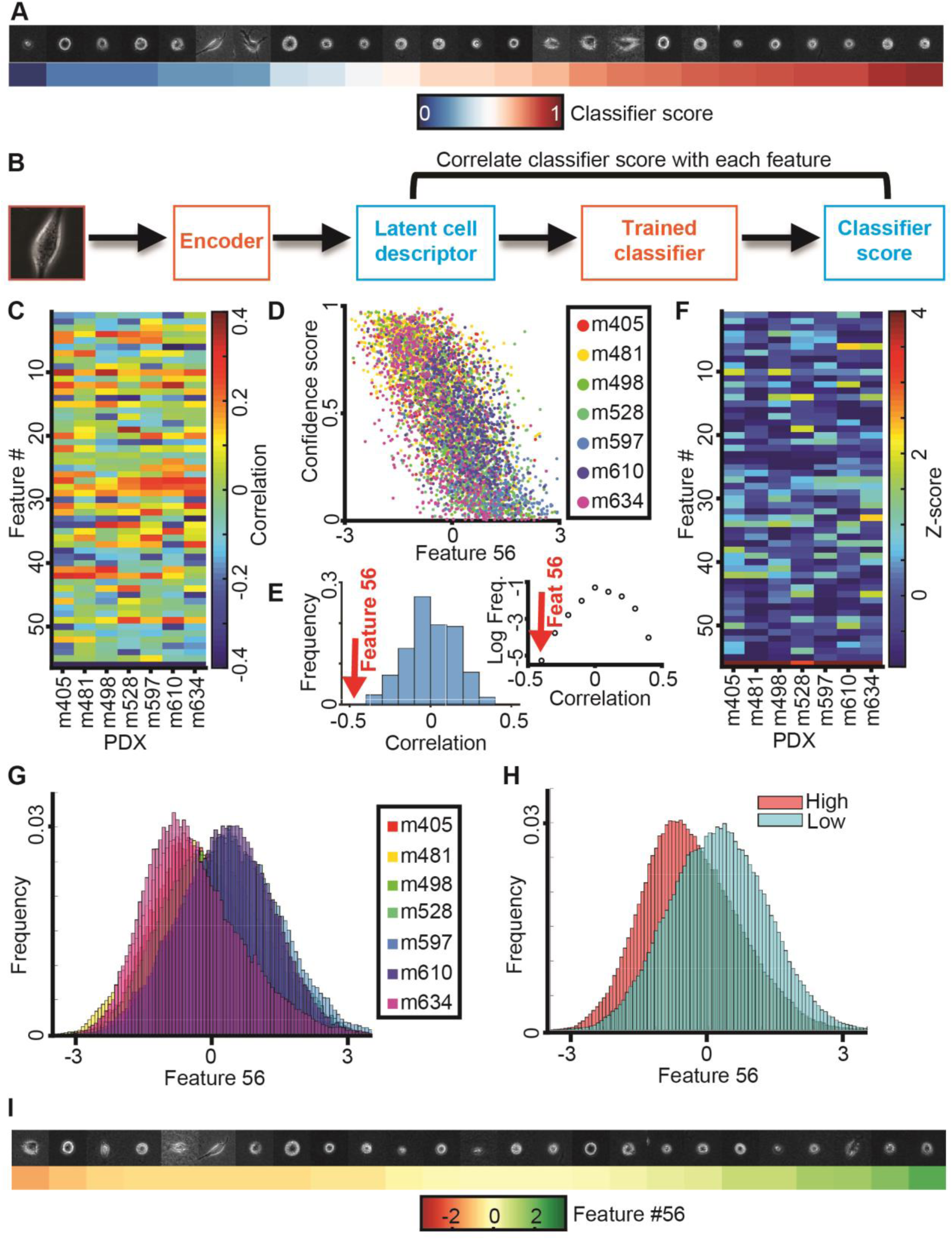
Metastatic efficiency is encoded by a single component of the latent space cell descriptor. (**A**) Gallery of snapshots of cells from a PDX (m610) ordered by their corresponding classifier score. (**B**) Approach: Each feature in the latent space cell descriptor is correlated with the score of the classifier trained to distinguish PDXs with high versus low metastatic efficiency. (**C**) Correlation between all 56 features (y-axis) and classifier scores for 7 PDXs (x-axis). (**D**) Value of feature #56 and classifier scores for individual cells color-grouped by PDX. (**E**) Distribution of the correlations from panel B; feature #56 (red arrow) is an obvious outlier. Left: distribution. Right: plot of log frequency for better visualization of feature #56. (**F**) Normalized correlation values (Z-scores) all 56 features (y-axis) and classifier scores (x-axis). Z-scores are calculated using the mean value and standard deviation of the distribution of correlation values in panel D. (**G**) Distribution of feature #56 values for cells grouped by association with a PDX. (**H**) Distribution of feature #56 values for cells grouped by association with low and high metastatic efficiency. (**I**) Gallery of snapshots of cells from PDX m610 in ascending order of the normalized value of feature #56. Note, high metastatic efficiency relates to negative, low metastatic efficiency to positive values of feature #56.

To probe which features encapsulated in the latent cell descriptor are most discriminative of the metastatic state we correlated each of the 56 features to the classifier score (Fig. 6B-C). The correlations were calculated independently for each PDX using a classifier blind to the PDX (see Fig. 4A). For all 7 PDXs the last feature #56 stood out as highly negatively correlated to the classifier scores (Fig. 6C-D). The correlation values fell significantly outside the range of correlations observed for any other feature (Fig. 6E-F). The distributions of feature #56 for individual PDXs clearly separated tumors with high versus low metastatic efficiency (Fig. 6G).

This finding was corroborated by a remarkable separation of the integrated distributions of high and low metastatic efficient PDXs (Fig. 6H). However, as with the classifier score (Fig. 6A), a series of random cell snapshots from one PDX in rank order of feature #56 values did not reveal a cell image pattern that could intuitively explain the meaning of this feature (Fig. 6I). This suggests that feature #56 encoded a multifaceted image property reflecting the metastatic potential of melanoma PDXs that cannot readily be grasped by visual inspection.

Intriguingly, when we applied the same feature-to-score correlation analysis to classifiers trained for discrimination of cell lines from PDXs, we found the three features #26, #27, and #36 as classification-driving (Fig. S16). These results highlight that for different classification tasks different feature subsets in the latent space cell descriptor capture distinguishing cell properties.

### Interpretation of classification-driving latent feature using generative models and time traces of feature values

Neither series of cell images rank-ordered by classification scores nor series rank-ordered by feature #56 offered a visual clue as to which image properties may determine a cell’s metastatic efficiency. We concluded that the natural variation of feature #56 values in our data was too low to give such clues and/or that the natural variation of features unrelated to metastatic efficiency largely masked image shifts related to the variation of feature #56 between PDXs with low and high metastatic efficiency. To glean some of the image properties that are controlled by feature #56 we exploited the network decoder to generate a series of “in silico” cell images in which, given a particular location of a cell in the latent space, feature #56 was gradually altered while fixing all other features (Fig. 7A). As expected, the changes in feature #56 negatively correlated with the changes they caused in the classifier score (Fig. 7B). The generative modeling brought two advantages over our previous attempts of visually interpreting feature #56: First, it allowed us to observe ‘pure’ image changes along a principal axis of metastatic efficiency change.

**Figure 7:**
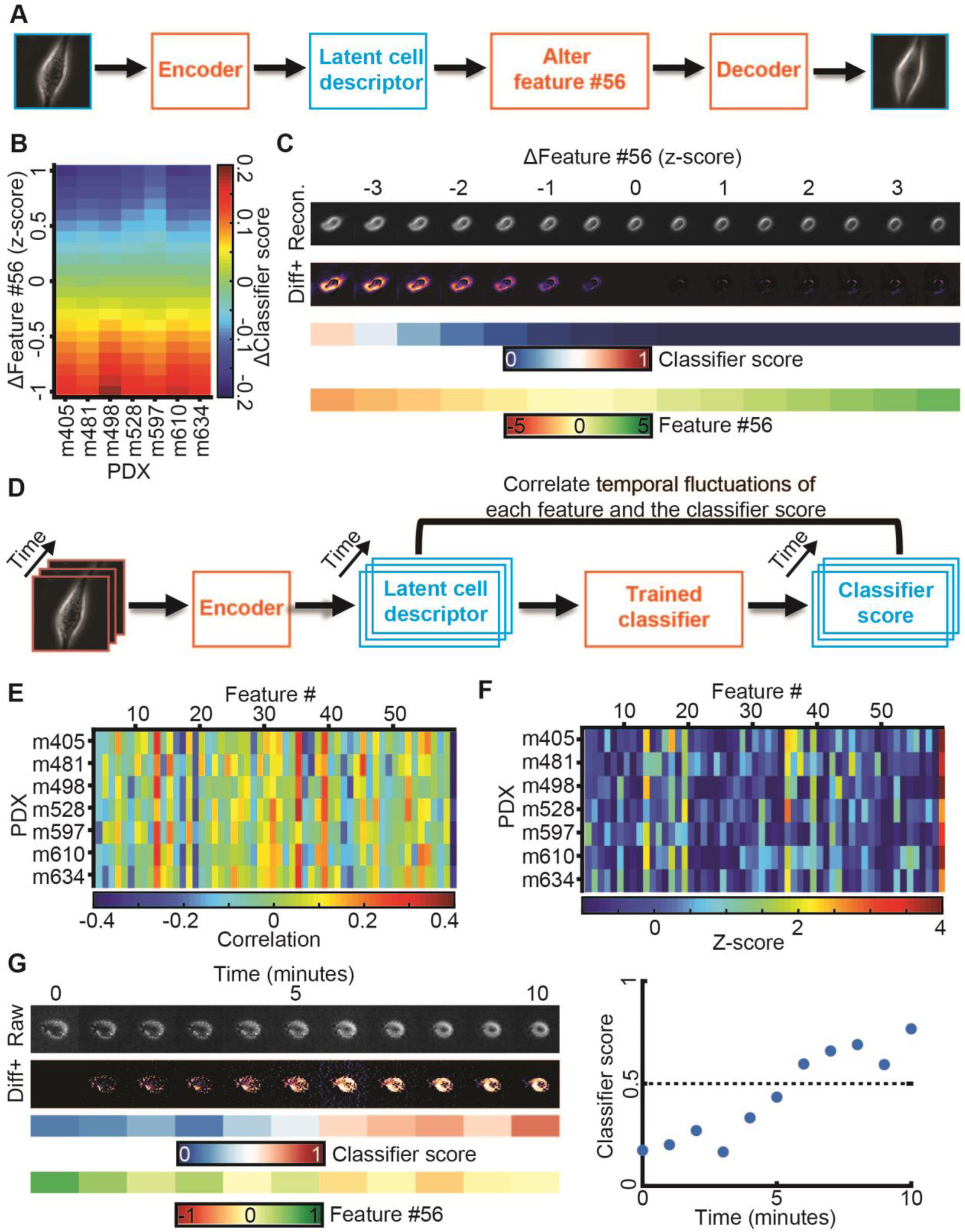
Generative modeling of cell images to interpret the meaning of feature #56. (**A**) Approach: alter feature #56 while fixing all other features in the latent space cell descriptor to identify interpretable cell image properties encoded by feature #56. (**B**) Shifts in feature #56 (y-axis, measured in z-score) negatively correlated with variation in the classifier scores. (**C**) *In silico* cells generated by decoding the latent cell descriptor of a representative m498 PDX cell under gradual shifts in feature #56 (“Recon.”). Visualization of the intensity differences between consecutive virtual cells (I_zscore_ - I_zscore+0.5_), only positive difference values are shown (“Diff+”). Changes in feature #56 are indicated in units of the z-score. The corresponding classifier’s score and value of feature #56 are shown. (**D**) Approach: correlating temporal fluctuations of each feature to fluctuations in the classifiers’ score. (**E**) Summary of correlations. Y-axis - different classifiers for each PDX. X-axis - features. Bin (x,y) records the Pearson correlation coefficients between temporal fluctuations in feature #x and the score of classifier #y over all cells of the PDX. (**F**) Normalization of correlation coefficients as a Z-score. Mean value and standard deviation are derived from the correlation values in panel E. (**G**) Following a m610 PDX cell spontaneously switching from the low to the high metastatic efficiency domain (as predicted by the classifier). Live imaging for 10 minutes. Left (top-to-bottom): raw cell image, diff+ images, classifier’s score, feature #56 values. Right: visualization of the classifier score as a function of time, switching from “low” to “high” in less than 10 minutes.

Second, it allowed us to shift the value of feature #56 significantly outside the value range of the natural distribution and thus to analyze the exaggerated cell images for emergent properties in cell appearance. Upon morphing a PDX cell classified as low metastatic within a normalized z-score range for feature #56 of [-3.5, 3.5], we observed two emergent properties in the high metastatic efficiency domain. The formation of pseudopodial extensions and changes in the level of cellular light scattering as observed by brighter image intensities at the cell periphery and interior (Fig. 7C). The pseudopodial activity was visually best appreciated when compiling the morphing sequences into videos that shift a cell classified as low metastatic towards the high metastatic efficiency domain (Video S5) and, vice versa, a cell classified as highly metastatic towards the low metastatic efficiency domain (Video S6).

Repeating the morphing for many PDX cells (Fig. S17, Video S7) underscores pseudopod formation and enhanced light scattering as the systematic and prominent factors that distinguish cells with low feature #56 values/high metastatic efficiency from those with high feature #56 values/low metastatic efficiency. Moreover, by variation of other features of the latent space cell descriptor we visually confirmed this combination of morphological properties was specifically controlled by feature #56 (Fig. S18).

To corroborate our conclusion from synthetic images we tested whether cells with significant plasticity in the classifier score displayed visually identifiable image transitions. First, we verified that temporal fluctuations in feature #56 negatively correlated with the temporal fluctuations in the classifier scores (Fig. 7D-F). Second, we confirmed that PDX cells spontaneously transitioning from a predicted low to a predicted high metastatic efficiency displayed increased light scattering (Fig. 7G, Video S8). We were not able to conclusively validate the enhanced protrusive activity in experimental data, perhaps due to the subtlety and subcellular localization of this phenotype.

### PDX-trained classifier can predict the metastatic potential of melanoma cell lines in mouse xenografts

Finally, we were interested in the capacity of PDX-trained classifiers to predict the metastatic outcome of any melanoma cell line that forms a tumor. We hypothesized that, despite the overall distinct morphologies of PDX and cell lines, the core differentiating properties between a low and high efficiency metastatic PDXs would be conserved for melanoma cell lines. Using the PDX-trained classifiers, A375, a BRAFV600E-mutated and NRAS wild-type melanoma cell line, which was originally excised from a primary malignant tumor (Davies et al., 2002; Ghandi et al., 2019; Giard et al., 1973; Kozlowski et al., 1984; Rozenberg et al., 2010; Tanami et al., 2004), was predicted as the most aggressive (Fig. 8A). MV3, a BRAF wild-type and NRAS-mutated melanoma cell line, that was originally excised from a metastatic lymph node and described as highly metastatic (Quax et al., 1991; Schrama et al., 2008; van Muijen et al., 1991), was predicted by the PDX-trained classifiers as the least aggressive (Fig. 8A). Consistent with our previous analyses of the influence of the latent space features on classification, feature #56 was lower for A375 than for MV3 (Fig. 8B). We subcutaneously injected luciferase-labeled versions of A375 and MV3 cells into the flanks of NSG mice (Methods). Over 25-34 days, both cell models formed robust primary tumors at the site of injection (Fig. 8C-D) as well as macrometastases in the lungs and in multiple other remote organs (Fig. 8E-F). Quantitative bioluminescence imaging of individual excised organs showed a significantly higher spreading to distant sites in A375 compared to MV3 cells (Fig. 8E-G). Intriguingly, primary tumors in MV3-injected mice grew much faster than in A375-injected mice (Fig. 8H), in contrast to being less aggressive in spreading to remote organs, suggesting that primary tumor growth is uncoupled from the ability to produce remote metastases (Ganesh et al., 2020; Quintana et al., 2012; Viceconte et al., 2017). Together, these data confirm that properties captured by feature #56 in the latent space cell descriptor define a specific gauge of the metastatic potential of melanoma that is independent of the tumorigenic potential.

**Figure 8:**
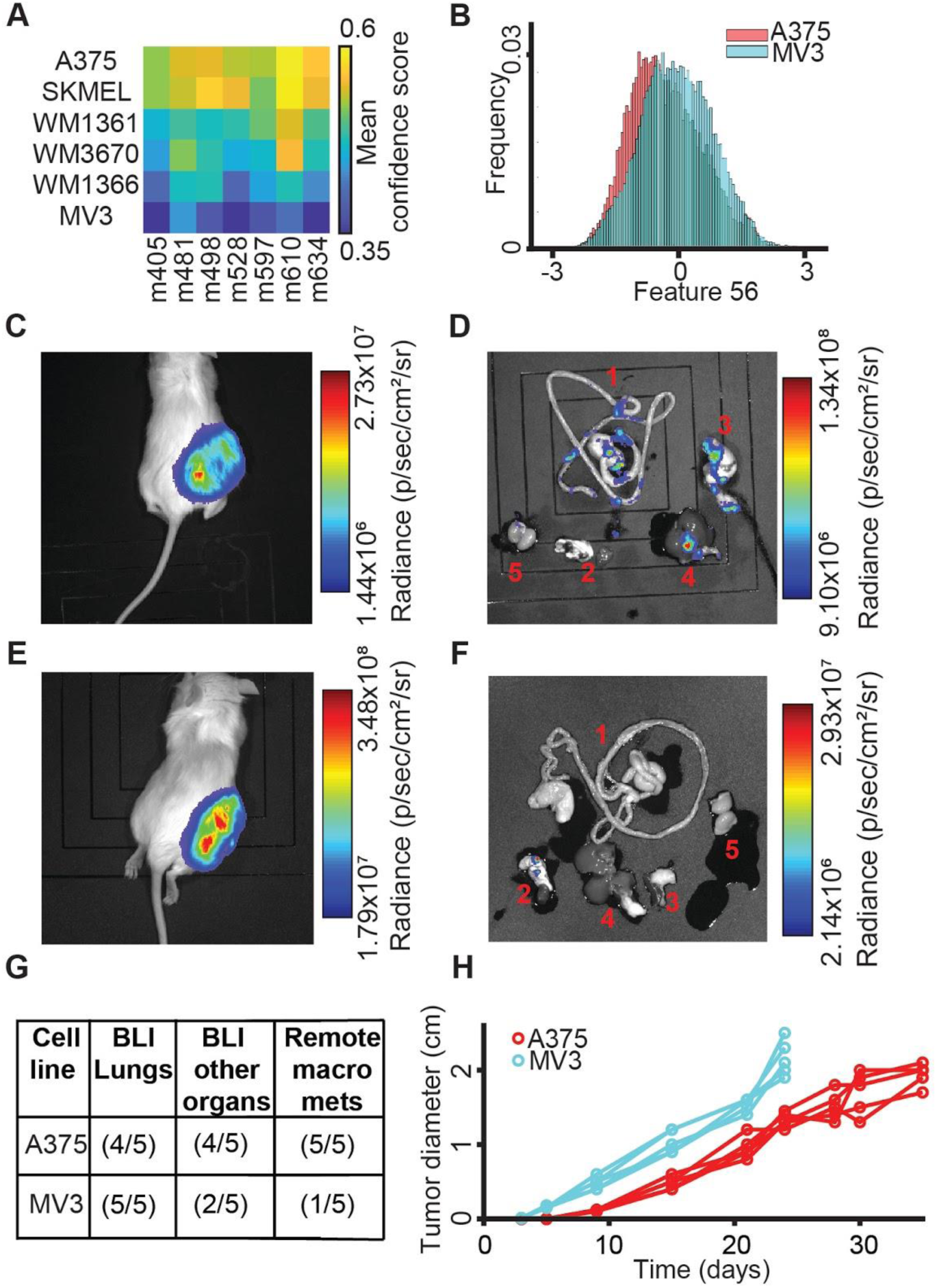
PDX-trained classifiers can predict the metastatic potential of melanoma cell lines in mouse xenografts. (**A**) All 7 PDX-trained classifiers consistently predicted that among the 6 analyzed cell lines A375 has the highest and MV3 the lowest metastatic efficiency. (**B**) The distribution of single cell values of feature #56 is lower for A375 than the distribution of values for MV3 cells. (**C, E**) BLI (Luminescence) of NSG mouse sacrificed 24-35 days after subcutaneous transplantation of 100 Luciferase-GFP+ cells from the A375 melanoma cell line (C) versus from the MV3 cell line (E). (**D, F**) Bioluminescence of organs dissected from the A375 xenografted mouse (D) and from the MV3-xenografted mouse (F). 1, Gastrointestinal Tract (GI); 2, Lungs and Heart; 3, Pancreas and Spleen; 4, Liver; 5, Kidneys and Adrenal glands. In the MV3, mouse metastases were mostly found in the Lungs. Black shades are mats on which the organs and mice are imaged (Methods). (**G**) Metastasis statistics from 5 mice for A375 and MV3 melanoma cell lines. (**H**) Primary tumors in MV3 mice grow faster than in A375 mice. Mice were sacrificed 24 days after injection with MV3, 35 days after injection with A375 cells. N = 5 mice for A375 and MV3 cell line. Statistics for tumor size after 24 days p-value = 0.0079 (Wilcoxon rank-sum test), fold = 1.6241.

## Discussion

### Visually unstructured properties of cell image appearance enable robust cell type classification

Morphology has long been a cue for cell biologists and pathologists to recognize cell type and abnormalities related to disease (Bakal et al., 2007; Chan, 2014; Eddy et al., 2018; Gordonov et al., 2015; Gurcan et al., 2009; López, 2013; Pavillon et al., 2018; Wu et al., 2020; Yin et al., 2013). In this study, we rely on the exquisite sensitivity of deep learned artificial neural networks in recognizing subtle but systematic image patterns to classify different cell types and cell states. To assess this potential we chose phase contrast light microscopy, an imaging modality that uses simple transmission of white or monochromatic light through an unlabeled cell specimen and thus minimizes experimental interference with the sensitive patient samples that we used in our study. A further advantage of phase contrast microscopy is that the imaging modality captures visually unstructured properties, which relate to a variety of cellular properties, including surface topography, organelle organization, cytoskeleton density and architecture, and interaction with fibrous extracellular matrix. We aimed to test whether these rich image properties would go beyond conventional descriptors of morphology in terms of distinguishing different cell types and states.

Our cell type classification rests on the combination of an unsupervised deep learned autoencoder and a conventional supervised classifier that discriminates between distinct cell categories. Autoencoders consist of two structurally symmetric networks, the first ‘encoding’ a full image into single numerical vector representative of the essential image content, which is referred to as the latent space feature vector; the second ‘decoding’ a feature vector in the latent space to synthesize a new image. In the classic autoencoding used in this study the encoding and decoding networks are trained to minimize the disagreement between the original and the synthetic image. Thus, autoencoding offers a powerful means for image denoising and data compression (Hinton and Salakhutdinov, 2006; Vincent et al., 2010). The generative capacity of the autoencoder architecture has recently also gained popularity for training mappings between different imaging modalities – in the context of microscopy to synthesize multi-spectral fluorescence images from bright-field images (Christiansen et al., 2018; Ounkomol et al., 2018), or super-resolution images from conventional fluorescence images(Ouyang et al., 2018; Weigert et al., 2018). In these cases, the decoders were trained such that they synthesized a new authentic target modality from the encoded latent space. These landmark studies have highlighted the unprecedented capacity of autoencoders to carry -- through the vehicle of a latent space representation -- visually hidden information from one imaging modality into visually accessible information in another imaging modality. By extracting the essence of a cell image, the latent cell descriptor is a low-dimensional representation potentially also suitable to distinguishing cell properties, i.e., two cells whose vectors fall in different locations of the latent space are expected to have different appearances. However, as the relation between the input image and the latent cell descriptor is governed by a highly nonlinear integration of convoluted image properties, the latent space is non-orthonormal and may contain discontinuities. While this characteristic of the latent space is less critical for mappings between identical or different imaging modalities, it precludes the application of any form of cell type classifier training under the assumption that proximity in latent space corresponds to proximity in image appearance. To impose this condition, it was necessary to constrain the encoder/decoder mappings by training an adversarial network in parallel to the decoder network (Fig. 2).

Our two-step implementation of unsupervised feature extraction and supervised classifier training allowed us to construct several different cell classifiers for different tasks using a one-time learned, common feature space. Specifically, we trained classifiers to distinguish melanoma cell lines from normal melanocytes, melanoma cell lines from expanded clones of these cell lines, melanoma cell lines from patient derived melanoma xenotransplants (PDX), and PDXs with high versus low metastatic efficiency (Figs. 4-5). All these tasks relied on the same fundamental capacity of a feature set to capture differences in cell appearance. Thus, the task of distinguishing melanoma cell lines from normal melanocytes could benefit from the information extracted from PDXs, while PDXs could be divided into groups with high versus low metastatic propensity with the support of information extracted from melanoma cell lines and untransformed melanocytes. Hence, sensitive classifiers could be trained on a relatively small data subsets – much smaller than would be required to train an ab initio deep-learned classifier for the same task. The approach is not only data-economical, but it greatly reduces computational costs as the deep learning procedure is performed only once on the full dataset. Indeed, in our study we learned a single latent feature space using time lapse sequences from over 12,000 cells (∼1.7 million snapshots); and then trained classifiers on data subsets that included labeled classes smaller than 1,000 cells. As an additional benefit of the orthogonalization of unsupervised feature extraction and supervised classifier training, we were able to evaluate the performance of our classifiers by repeated leave-one-out validation, verifying that the discriminative model training is completely independent of the cell type at test. A similar evaluation strategy requiring the repeated re-training of a deep learned classifier would likely become computationally prohibitive. Future studies will illuminate the robustness of our melanoma latent representation to similar classification tasks on other cancers or entirely unrelated cell type distinctions.

### Application of cell type classification to the prediction of metastatic efficiency

Among multiple cell type classification tasks, we were able to distinguish the metastatic efficiency of stage III melanoma harvested from a xenotransplantation assay that had previously been shown to maintain the patient outcome (Quintana et al., 2012). Therefore, the proposed classification scheme opens the window to a potential clinical lab test based on live cell imaging for patients presenting with metastatic spreading to the lymph system. Accordingly, we refer to our approach as Live Cell Histology.

Our classifier perfectly distinguished PDX populations that have shown high versus low metastatic spread in patients. At the single cell level the classifier accuracy dropped to 70%. This is not necessarily a weakness of the classifier but speaks to the fact that tumor cell populations grown from a single cell clone are not homogeneous in function and/or appearance. Importantly, our estimates of classifier accuracy relies on leave-one-out strategies where the training set and the test set were completely non-overlapping, both with regards to the classified cell type and to the days the classified cell type was imaged. Thus, it can be assumed that the reported accuracies can be reproduced on new, independent PDX models.

Besides numerical testing, we validated the accuracy of our classifiers high versus low metastatic efficiency in a fully orthogonal experiment. We applied the PDX-trained classifiers to predict the metastatic efficiency of well-established melanoma cell lines and validated their predictions in mouse xenografts. We emphasize that the PDX-trained classifier has never encountered a cell line and that despite the significant differences between cell lines and PDXs (Fig. 4H-J), the classifier correctly predicted high metastatic potential for the cell line A375 and low potential for MV3 (Fig. 8). Intriguingly, the aggressiveness in primary tumor growth was reverted between the two cell lines, in agreement with many other reports suggesting that tumorigenesis and metastasis are unrelated phenomena (Ganesh et al., 2020; Quintana et al., 2012; Viceconte et al., 2017). This shows that the latent feature space encodes cell properties that specifically contribute to cell functions required for metastatic spreading and that these features are orthogonal to features, which allow the training of a classifier distinguishing cell lines from patient-derived melanoma models.

### Image-based classifiers are more predictive of metastatic potential than the mutational profile

The development of the live cell histology pipeline into a clinical test for metastatic risk assessment would complement the current arsenal of tools for molecular diagnostics in precision medicine, including genomic profiling, with a predictor how aggressive a patient with early stage metastasis should be treated. While metastatic melanoma are expected to harbor a ‘standard’ set of primary mutations, such as those in BRAF or NRAS (Jakob et al., 2012) – and indeed all our PDX models and metastatic cell lines do contain an activating mutation in either one of these genes (Table S2) – we were curious as to whether secondary mutations in the genomic profiles of these cell models would encode information on the metastatic propensity, similar to the deep-learned latent feature vector derived from live cell microscopy, or whether genomics and imaging offered largely complementary data. To address this question we scrutinized the distributions of genomic distances among the PDX cell models and two cell lines vis-à-vis the distance distributions in the latent feature space. The conclusion from these experiments was that the states of oncogenic/likely-oncogenic mutations in the 20 most mutated genes in melanoma (Hodis et al., 2012) were insufficient for a prediction of the metastatic efficiency (Fig. S19). In fact, the oncogenic/likely-oncogenic mutations in the genes were not more predictive than non-oncogenic mutations or an unbiased analysis of a full panel of 1400 genes for metastatic states. Thus, for specific functional analyses, image-based classifiers are indeed a promising complement to profiles of genomic variants.

While the disconnect between mutational patterns and metastatic efficiency may be specific to the notoriously heterogeneous genomes of metastatic melanoma (Harbst et al., 2014), these analyses highlight the value of designing assays in cancer diagnosis that are close to the actual cancer cell function. As illustrated in Fig. S20, the link between the genomic cell state and metastasis-enabling cell functions progresses through several states, each of which is perturbed by uncontrolled variation and feedbacks that further disconnect the genome from cell function. Most importantly, the link is further weakened by influences from cell environmental variables. Thus, our experiments confirm the validity of the century-old paradigm of pathology that asserts close proximity of cell image appearance and disease-driving cell function, especially in multi-factorial diseases like cancer.

### Interpretation of latent features discriminating high and low metastatic cell propensity

Deep Learning Artificial Neural Networks have revolutionized machine learning and computer vision as powerful tools for complex pattern recognition. However, the often cited weakness of these techniques is the lack of an intuitive explanation of which parts of the data are particularly meaningful in defining the extracted pattern. While in some applications, such as image segmentation, image restoration or mapping between imaging modalities, a well-validated outcome of a network has been satisfactory (Christiansen et al., 2018; Fang et al., 2019b; Guo et al., 2019; Hershko et al., 2019; Hollandi et al., 2019; LaChance and Cohen, 2020; Moen et al., 2019; Nehme et al., 2018; Ounkomol et al., 2018; Ouyang et al., 2018; Rivenson et al., 2019; Wang et al., 2019; Weigert et al., 2018; Wu et al., 2019), there is increasing mistrust in results produced by ‘black-box’ neural networks. Aside from increasing the confidence, the analysis of the properties – also referred to as ‘mechanisms’ – of the pattern recognition process can potentially generate insight of a biological/physical phenomenon that escapes the analysis driven by human intuition. Thus, while our classification results clearly demonstrated the predictive value of the deep-learned latent representation of cell appearance, we were wondering whether we could extract information about some of the key physical attributes that permitted our classifiers to discriminate cells with high versus low metastatic propensity. This would allow the formulation of hypotheses about hallmark properties of metastatic cells.

Recent advances in medical imaging explicitly identified local image sub-regions that determine the training of classifier deep neural networks (Courtiol et al., 2019; Fu et al., 2019; Pan et al., 2019; Shamai et al., 2019). Localization of sub-regions that were particularly important for the classifier result permitted a visual assessment and pathological interpretation of distinctive image properties. Such approaches are not suitable in our case where the driver of the classification was a complex integration of multiple co-localized image properties rather than an image sub-region. Because of the orthogonalization of feature space construction and classifier training we could elegantly tackle the quest for interpretability and extract visual cues for inspection of the classifier-relevant cell appearances. By exploiting the single cell variation of the latent feature space occupancy and the associated variation in the scoring of a classifier discriminating high from low metastatic melanoma, we identified feature #56 as predominant in prescribing metastatic propensity. Of note, the feature-to-classifier variation analysis is not restricted to determining a single discriminatory feature, as illustrated in our study with the example of multiple features driving the discrimination between cell lines and PDXs (Fig. S16). Thus, other applications of feature-to-classifier variation analysis may require more complex strategies to identify mixtures of latent space components that encode the information required for the discrimination of a specific cell function. Visual inspection of cell images ranked by the classifier score or feature #56 did not reveal any salient cell image appearance that would distinguish efficiently from inefficiently metastasizing cells (Fig. 6A,I). These particular image properties were masked by the vast variability of cell appearances that are unrelated to the metastatic function. Moreover, the function-driving feature #56 may be the nonlinear combination of multiple image properties and thus not discernible to the observer’s eye.

To test whether feature #56 encodes image properties that are human-interpretable but buried in the intrinsic heterogeneity of cell image appearances, we exploited the generative power of our autoencoder. While the encoder training optimized a minimal space to represent cell-to-cell variation in the raw images, the decoder was trained to generate realistic in silico cell images from this minimal space. Thus, we could ‘shift’ cells around in the latent feature space and observe the associated shifts in cell appearance. We thus could examine how cell appearances would change as the values of feature #56 exceeded the natural range of the feature in our experimental data, while fixing the other 55 feature values. Hence, the combination of exaggeration and purity allowed us to generate human discernible changes in image appearance that correspond to a shift in metastatic efficiency.

Once we had an idea of what to look for from exaggerated in silico images, we could validate the predicted appearance shifts in experimental data. We searched our data set for cells whose classification score and feature #56 values drifted from a low to high metastatic state or vice versa. We supposed that during such spontaneous dynamic events the variation in cell image appearances were, for a brief time window, dominated by the variation in feature #56 whereas other features influenced the image only marginally. Therefore, time-resolved data may present transitions in cell image appearance comparable to those induced by selective manipulation of latent space values along the direction of feature #56. It is highly unlikely to find a similarly pure transition between any pair of cells, explaining why we were unable to discern differences between cells with low and high metastatic efficiency in feature #56 ordered cell image series.

Even though the time-resolved amplification of feature #56 was weaker, and the fluctuations of other features were greater than the controlled shifts of in silico generated cells, we were able to verify the discriminating cellular properties by isolating and focusing on the specific in silico generated visual hypotheses.

Analyses of appearance shifts in both exaggerated in silico images and selected experimental images unveiled two functional hallmark properties of highly metastatic melanoma cells. First, these cells seemed to form pseudopodial extensions, especially in image simulations of the stereotypical transitions between low and high metastatic cell states (Fig. 7C, Fig. S17, Video S5, Video S6). Because of its subtlety, this phenotype was more difficult to discern visually during spontaneous transitions (Fig. 7G). Second, images of cells in a highly metastatic state displayed brighter cell peripheral and interior signals, indicative of alteration in cellular light scattering. Because light scattering affects the image signal globally, this phenotype was clearly apparent in simulations (Fig. 7C, Fig. S17, Video S5, Video S6) and experimental time lapse sequences of transitions in metastatic efficiency (Fig. 7G, Video S8). Importantly, neither one of the two phenotypes follows a mathematically intuitive formalism that could be implemented as an ab initio feature detector. This highlights the power of deep learned networks in extracting complex cell function-driving image appearances.

Pseudopodial extensions play critical roles in cell invasion and migration. However, at least in a simplified migration assay in tissue culture dishes, the highly metastatic cell population did not exhibit significantly enhanced migration (Fig. S14). Recent work has suggested mechanistic links between enhanced branched actin formation in pseudopdial cell compartments and enhanced cell cycle progression (Mohan et al., 2019; Molinie et al., 2019), especially in micro-metastases. Therefore, we lean towards an interpretation that connects the predicted metastatic efficiency under pseudopod formation to increased proliferation and survival.

The observation that light scattering can indicate metastatic efficiency suggests that the cellular organelles and processes captured by light scattering are relevant to the metastatic process (Schürmann et al., 2015). Indeed, differences in light scattering upon acetic acid treatment are often used to detect cancerous cells in patients (Marina et al., 2012). Although the mechanisms underlying light scattering of cells are unclear, intracellular organelles such as phase separated droplets (Falke et al., 2019) or lysosomes will be detected by changes to light scattering (Choi et al., 2007). With the establishment of our machine-learning based classifier, we are set to systematically probe the intersection of hypothetical metastasis-driving molecular processes, actual metastatic efficiency, and cell image appearance in follow-up studies.

## Methods

### Patient-derived xenograft (PDX) melanoma cells

Populations of primary melanoma cells were created from tumors grown in murine xenograft models as described previously (Quintana et al., 2010). Briefly, cells were suspended in Leibovitz’s L-15 Medium (ThermoFisher) containing mg/ml bovine serum albumin, 1% penicillin/streptomycin, 10 mM HEPES and 25% high protein Matrigel (product 354248; BD Biosciences). Subcutaneous injections of human melanoma cells were performed in the flank of NOD.CB17-*Prkdc*^scid^ *Il2rg^tm1Wjl^*/SzJ (NSG) mice (Jackson Laboratory). These experiments were performed according to protocols approved by the animal use committees at the University of Texas Southwestern Medical Center (protocol 2011-0118). After surgical removal, tumors were mechanically dissociated and subjected to enzymatic digestion for 20 min with 200 U ml−1 collagenase IV (Worthington), 5 mM CaCl2, and 50 U ml−1 DNase at 37oC. Cells were filtered through a 40 μm cell strainer to break up cell clumps and washed through the strainer to remove cells from large tissue pieces.

### Cell culture and origin

Cell cultures were grown on polystyrene tissue culture dishes to confluence at 37°C and 5% CO2. Melanoma cells derived from murine PDX models were gifts from Sean Morrison (UT Southwestern Medical Center, Dallas, TX) and cultured in medium containing the Melanocyte Growth Kit and Dermal Cell Basal Medium from ATCC. Primary melanocytes were obtained from ATCC (PCS-200-013) and grown in medium containing the Melanocyte Growth Kit and Dermal Cell Basal Medium from ATCC. The m116 melanocytes, a gift from J. Shay (UT Southwestern Medical Center, Dallas), were derived from fetal foreskin and were cultured in medium 254 (Fisher). A375 cells were obtained from ATCC (CRL-1619). SK-Mel2 cells were obtained from ATCC (HTB-68). MV3 cells were a gift from Peter Friedl (MD Anderson Cancer Center, Houston, TX). MV3 and A375 cells were cultured in DMEM with 10% FBS. WM3670, WM1361, and WM1366 were obtained directly from the Wistar Institute and cultured in the recommended medium (80% MCDB1653, 20%, 2% FBS, CaCl2 and bovine insulin).

### PDX-derived cell culture

We found that melanoma cell cultures derived from PDX tumors exhibited variable responses to traditional cell culture practices. Although some of the cell cultures retained high viability and proliferated readily, others exhibited significant cell death and failed to proliferate. We determined that frequent media changes (<24 hrs) and subculturing only at high (>50%) confluence dramatically increased the viability and proliferation of PDX-derived cell cultures.

Although we observed no correlation between metastatic efficiency and robustness in cell culture, we followed these general cell culture practices for all PDX-derived cultures.

### Clonal cell line experiments

To create cell populations “cloned” from a single cell, cells were released from the culture dish via trypsinization and passed through a cell strainer (Fischer; 07-201-430) to ensure single-cell solution, counted and then seeded on a 10 cm polystyrene tissue culture dish at low density of 350,000 cells/10 ml of phenol-red free DMEM. Single cells were identified via phase-contrast microscopy. The single cells were isolated using cloning rings (Sigma; C1059) and expanded within the ring. For clonal medium changes, the medium was aspirated within the cloning rings. Subsequently, conditioned medium from a culture dish with corresponding confluent cells were passed through a filter (Fischer; 568-0020), which removed any cells and cell debris and then added to each cloning ring. Once confluent within the cloning ring, the clonal populations were released via trypsinization inside the cloning ring, transferred to individual cell culture dishes, and allowed to expand until confluence.

### Bioluminescence imaging of NSG mice with melanoma cell lines

Injection of melanoma cells, monitoring of mice, dissection of mice, and imaging were all done as described in Quintana & Piskounova et al. (Quintana et al., 2012). Briefly, 100 Luciferase-GFP+ cells were injected into the right flank. Mice were monitored until the tumor at the site of injection reached 2 cm in diameter. Mice injected with MV3 were sacrificed 24 days after injection and A375 sacrificed 35 days after injection. The stomach, gut, rectum, and esophagus were labeled as the gastrointestinal tract. The black shades are mats that were used to image the mice’s organs. Some mouse/organ images have mats with (Fig. 8D) and without (Fig. 8F) gridlines.

### Quantification of metastatic efficiency in NSG mice

We used three measures to assess metastatic efficiency (Quintana et al., 2012). First, detection of BLI in the lungs. Second, detection of BLI in multiple organs beyond the lungs. Third, identification of “visceral metastasis”, macrometastases visually identifiable without BLI, see details in (Quintana et al., 2012). Macrometastases without a BLI signal occurred exclusively in remote organs.

### Measuring extracellular acidification rate, oxygen consumption rate and proliferation rate in PDXs

Cells were trypsinized and counted, and 4.0 x 104 cells/mL of cell suspension was added to Seahorse XFe24 Cell Culture Microplates (Agilent; 100777-004). The Seahorse culture plate was then placed in a 37°C incubator overnight. The Seahorse Extraflux Assay Kit was prepared as follows. Each well of a Seahorse XFe24 Utility Plate was filled with 200 μL of Seahorse XF Calibrant Medium (Agilent; 100840-000). The Seahorse sensor cartridge was added to the utility plate, submerging the sensors in the Calibrant Medium to hydrate. The Extraflux Assay kit was then placed in a CO2 free, 37°C incubator overnight. The following day, cells were washed with 1X PBS. Then, 200 μL XF Calibrant Medium was gently added to each well. Cells were inspected to verify that washing and medium additions did not detach or disturb cells. Then, cells were left in a CO2 Free, 37°C incubator for one hour to normalize pH. Finally, the Seahorse sensor cartridge was placed in the XFe24 culture plate and moved into the Agilent XFe24 Analyzer.

We used the Click-iT EDU based assay (ThermoFisher; C10340) to measure cell proliferation rate as described previously (Murali et al., 2019). For all proliferation measurements, melanoma cells were plated in a 24-well plate at a confluence of 4.0 x 104 cells/well and incubated at 37°C overnight. The next morning, the media was drawn off and replaced with MGM medium containing a 20 μM concentration of EDU. Cells were incubated at 37°C in the EDU containing medium for 24 hours. Following this incubation, cells were fixed in 4% PFA for 10 minutes and washed twice with 1X PBS. Fixed cells were then permeabilized with 0.5% tritonX 100 (FisherBioreagents; BP151-500) in PBS for 20 minutes and washed twice with 1X PBS. Cells were then stained with Alexa Flour 647 Azide at a 1:100 dilution in DMSO for 30 minutes. Cells were then washed three times with 1X PBS. After EDU labeling, cells were also labeled with DAPI at 1:1000 dilution in PBS and for 5 minutes. Cells were then washed four times with 1X PBS and imaged at 20x magnification on a Nikon Eclipse Ti live cell microscope. Proliferation was quantified in these images using a point source detection algorithm described previously (Mohan et al., 2019).

### Targeted sequencing cancer-related genes and copy number variation analysis

Targeted sequencing of exons of 1385 cancer-related genes was performed by the Genomics and Molecular Pathology Core at UT Southwestern Medical Center as previously described (Zhang et al., 2020). Sequencing was performed on 6 out of 7 PDXs and the two cell lines A375 and MV3. Due to the difficulty in expanding the cells of PDX m528 in culture, we were not able to sequence this PDX. From the raw variant calling files, high confidence variants were determined by filtering variants found to have (a) strand bias, (b) depth of coverage < 20 reads and alt allele frequency < 20%. Common variants were filtered if they were in > 1% allele frequency in any population (Karczewski et al., 2020). Oncogenic potential was assess using oncokb-annotator (https://github.com/oncokb/oncokb-annotator). Summary tables of high-confidence variants of melanoma PDXs and cell lines were assembled in Table S2.

### Live cell imaging

Live cell phase contrast imaging was performed on a Nikon Ti microscope equipped with an environmental chamber held at 37oC and 5% CO2 in 20x magnification (pixel size of 0.325μm). In order to prevent morphological homogenization and to better mimic the collagenous ECM of the dermal stroma, we imaged cells on top of a thick slab of collagen. Collagen slabs were made from rat tail collagen Type 1 (Corning; 354249) at a final concentration of 3 mg/mL, created by mixing with the appropriate volume of 10x PBS and water and neutralized with 1N NaOH. A total of 200 μL of collagen solution was added to the glass bottom portion of a Gamma Irradiated 35MM Glass Bottom Culture Dish (MatTek P35G-0-20-C). The dish was then placed in an incubator at 37°C for 15 minutes to allow for polymerization.

Cells were seeded on top of the collagen slab at a final cell count of 5000 cells in 400 uL of medium per dish. This solution was carefully laid on top of the collagen slab, making sure not to disturb the collagen or spill any medium off of the collagen and onto the plastic of the MatTek dish. The dish was then placed in a 37°C incubator for 4 hours. Following incubation, one mL of medium was gently added to the dish. The medium was gently stirred to suspend debris and unattached cells. The medium was then drawn off and gently replaced with two mL of fresh medium.

### Single cell detection and tracking

We took advantage of the observation that image regions associated with “cellular foreground” had lower temporal correlation than the background regions associated with the collagen slab because of their textured and dynamic nature. This allowed us to develop an image analysis pipeline that detected and tracked cells without segmenting the cell outline. This approach allowed us to deal with the vast variability in the appearance of the different cell models and batch imaging artifacts in the phase-contrast images. The detection was performed in super-pixels with a size equivalent to a 10 x 10 μm patch. For each patch in every image, we recorded two measurements, one temporal- and the other intensity-dependent (see details later), generating two corresponding downsampled images reflecting the local probability of a cell being present.

We used these as input to a particle tracking software, which detected and tracked local maxima of particularly high probability (Aguet et al., 2013). The first measurement captures the patch’s maximal spatial cross-correlation from frame *t* to frame *t+1* within a search radius that can capture cell motion up to 60 μm/hour. The second measurement used the mean patch intensity in the raw image to capture the slightly brighter intensity of cells in relation to the background in phase-contrast imaging. Notably, our reduced resolution in the segmentation-free detection and tracking approach would break for imaging in higher cell densities. A bounding box of 70 x 70 µm around each cell was defined and used for single cell segmentation and feature extraction (details will follow). We excluded cells within 70µm from the image boundaries to avoid analyzing cells entering or leaving the field of view and to avoid the characteristic uneven illumination in these regions.

### Single cell segmentation in phase-contrast imaging and shape feature extraction

Label-free cell segmentation is a challenging task, especially in the diverse landscape of shapes and appearance of the different melanoma cell systems we used. We used the LEVER (Winter et al., 2016) (downloaded from https://git-bioimage.coe.drexel.edu/opensource/lever), a designated phase-contrast cell segmentation algorithm to segment single cells within the bounding boxes identified by the previously described segmentation-free cell tracking. Briefly, the LEVER segmentation is based on minimum cross entropy thresholding and additional post-processing. While the segmentation was not perfect, it generally performed robustly to cells from different origins and varied imaging conditions (Fig. S11). We used MATLAB’s function *regionprops* to extract 13 standard shape features from the segmentation masks produced by LEVER. These included: Area, MajorAxisLength, MinorAxisLength, Eccentricity, Orientation, ConvexArea, FilledArea, EulerNumber, EquivDiameter, Solidity, Extent, Perimeter, PerimeterOld.

### Unsupervised feature extraction with Adversarial Autoencoders

We have developed an unsupervised, generative representation for capturing cell image features using Adversarial Autoencoders (AAE) (Goodfellow et al., 2014; Makhzani et al., 2015). The autoencoder learns a compressed representation of cell images by encoding the images using a series of convolution and pooling layers leading ultimately to a lower dimensional embedding, or latent space. Points in the embedding space can then be decoded by a symmetric series of layers flowing in the opposite direction to reconstruct an image that, once trained, ideally appears nearly identical to the original input (Hinton et al., 2006). The training/optimization of the AAE is regularized (by using a GAN during training) such that points close together in the embedding space will generate images sharing close visual resemblance/features (Makhzani et al., 2015).

This convenient property can also generate synthetic/imaginary cell images to interpolate the appearance of cells from different regions of the space. We used the architecture from Johnson et al. (Johnson et al., 2017), that was based on the network presented in (Makhzani et al., 2015).

Johnson’s network includes an AAE that learns to reconstruct landmarks of the cell nucleus and cytoplasm. The adversarial component teaches the network to discriminate between features derived from real cells and those drawn randomly from the latent space. We trained the regularized AAE with bounding boxes of phase-contrast single cell images (of size 70µm x 70 µm, or 217 x 217 pixels) that were rescaled to 256×256 pixels. The network was trained to extract a 56-dimensional image encoding representation of cell appearance. This representation and its variation over time were used as descriptors for cell appearance and action. We adapted Torch code from https://github.com/AllenCellModeling/torch_integrated_cell (Arulkumaran, 2017; Johnson et al., 2017) for unsupervised AAEs, and adjusted it to execute on our high-performance computing cluster. Torch (Collobert et al., 2011) is a Lua script-based scientific computing framework oriented towards machine learning algorithms with underlying C/CUDA implementation.

### Encoding temporal information

We compared three different approaches to incorporating temporal information when using either the autoencoder-based representation or the shape-based representation of cell appearance. First, static snapshot images ignoring the temporal information. Second, averaging the cell static descriptors along a cell’s trajectory, canceling noise for cells that do not undergo dramatic changes. Notably, the resulting cell descriptor matches the static descriptor in size and features.

Accordingly, classifiers that were trained on average temporal descriptors could be applied to static snapshot descriptors (see Figs. 6-7). In the third encoding we relied on the ‘bag of words’ approach (Sivic and Zisserman, 2009), in which each trajectory is represented by the distribution of discrete cell states, termed ‘code words’. A ‘dictionary’ of 100 code words was predetermined by k-means clustering (MacQueen, 1967) on the full dataset of cell descriptors.

### Dimensionality reduction

We used tSNE (Fig. 3C) and PCA (Fig. S5) for dimensionality reduction. Each cell was represented by its time-averaged descriptors in the latent space. For tSNE we used a GPU-accelerated implementation, https://github.com/CannyLab/tsne-cuda (Chan et al., 2018).

### Discrimination analysis

We used Matlab’s vanilla implementation of Linear Discriminant Analysis (LDA) for the discrimination tasks (Figs. 4-5) and to identify the cellular phenotypes that correlate with low or high metastatic efficiency (Figs. 6-7). The feature vector for each cell was given by a the normalized latent cell descriptor extracted by the autoencoder. Normalization of each latent cell descriptor component to a z-score feature was accomplished as follows. The mean (μ) and standard deviation (σ) of a latent cell descriptor component were calculated across the full data set of cropped cell images and used to calculate the corresponding z-score measure: x^norm^= (x−μ)/σ, i.e., the variation from the mean values in units of standard deviation that can later be compared across different features.

For each classification task, the training data was kept completely separate from the testing data. Training and testing sets were assigned according to two methodologies. First, hold out all data from one cell type and train the classifier using all other cell types (Fig. 4A). Second, hold out all data from one cell type imaged in one day as the test set (“cell type - day”, e.g., Fig. 5F) and train the classifier on all other cell types excluding the data imaged on the same day as the test set (Fig. S6). This second approach trained models that had never seen the cell type or data imaged on the same day of testing. In both classification settings we balanced the instances from each class for training by randomly selecting an equal number of observations from each class. This scheme was used for classification tasks involving class set labels containing more than one cell type: cell lines versus melanocytes, cell lines versus clonally expanded cell lines, cell lines versus PDXs, low versus high metastatic efficiency in PDXs (Figs. 4-5). For statistical analysis, all the cells in a single test set are considered as a single independent observation. Hence, “cell type - day” testing sets provide more independent observations (N) at the cost of fewer cells imaged in each day compared to testing set of the form of “cell type”.

We used bootstrapping to statistically test the ability to predict metastatic efficiency from samples of 20 random cells. This was performed for “cell type” (Fig. 5D) or “cell type - day” (Fig. 5G) test sets. For each test set, we generated 1000 observations by repeatedly selecting 20 random cells (with repetitions), recorded the fraction of these cells that were classified as low efficiency and the 95% confidence interval of the median. Statistical significance in all settings was inferred using two statistical tests using each test set classifier’s mean score: (1) The nonparametric Wilcoxon signed-rank test, considering the null hypothesis that the classifiers scores of observations from the two classes stem from the same distribution; (2) The Binomial test, considering the null hypothesis that the classifier prediction is random in respect to the ground truth labels. For inference of phenotypes that correlate with metastatic efficiency (Fig. 7) we used the classifier that was trained on the mean latent cell description along its trajectory (which proved to be superior to training with single snapshots) on latent cell descriptors derived from single snapshots, which hold the same, just noisier features.

The area under the Receiver Operating Characteristic (ROC) curve was recorded to assess and compare the discriminative accuracy of different tasks (Figs. 4-5). The true-positive rate (TPR) or sensitivity is the percentage of “low” metastatic cells classified correctly. The false-positive rate (FPR) or (1-specificity) is the percent of “high” metastatic cells incorrectly classified as “low”. Area under the ROC curve (AUC) was used as a measure of discrimination power. Note that the scores of all cells from all relevant cell types were pooled together for this analysis.

Importantly, different classifiers can produce different scores, which means that our analysis provides a lower bound (pessimistic estimation). ROC analysis could not be applied for individual (held-out) test sets because they consist of only a single ground truth label. We used the web-application PlotsOfData (Postma and Goedhart, 2019) to generate all boxplots.

### Measuring cell plasticity

We defined a “plastic” or “transition” event for a PDX cell when the classifier predicted a switch from high-to-low or low-to-high according to the following criteria: (1) Significant change of at least 0.2 in classification score; (2) Maintaining the same label for at least 5 frames (5 minutes) before and after switching labels. Importantly, the PDX from which a cell was taken has one known metastatic efficiency label. However, the classifier is not aware of that label (and is not perfect) and can predict transitions between the “low” and “high” states in the same cell. For each PDX, we measured the average rate of transition over all cells and the fraction of cells that underwent such transitions along their time-lapse imaging.

### Correlating classifier scores to genomic mutation markers

We calculated a distance matrix to assess the similarity between all pairs of PDXs and the cell lines A375 and MV3. The distances were calculated in terms of the classifier score and of genomic mutation panels. m528 was excluded from the analysis due missing sequencing data (see above). For the distance matrix of the classifier score, we calculated the Jensen-Shannon (JS) divergence (Lin, 1991) between the distributions of single cell classifier scores using the corresponding PDX-based classifiers (see discrimination analysis section in the Methods). For the cell lines, a new classifier was trained using all cells from all seven PDXs. This classifier was used to determine the classifier score for A375 and MV3. For each cell type, the distribution was approximated with a 25 bin histogram. JS divergence was calculated on pairs of cell type classifier score distributions.

To calculate distance matrices based genomic mutations we considered three panels of established melanoma genomic mutation markers. Two genomic mutation panels were derived from variation of exomes associated with 1385 cancer-related genes (see above). Mutations in commonly mutated genes in melanoma (Hodis et al., 2012) were annotated using OncoKB (Chakravarty et al., 2017) and divided into (i) oncogenic or likely oncogenic (Table S3, Fig. S19B) and (ii) benign or unannotated (“non-oncogenic”) (Table S4, Fig. S19C). Mutational based genetic distances were derived by converting mutation scores to a binary state (1=presence, 0=absence) and computing the Jaccard index (Jaccard, 1912) between cell types. In Fig. S19D we calculated distances using MASH (Ondov et al., 2016), which compared the K-mer profiles between samples, thus giving a distance of the raw sequence data, without biases introduced in the alignment and variant calling analysis.

The distance matrices derived from classifier scores and mutational states were correlated (Pearson correlation) to assess whether the genomic mutation state and image-derived classifier scores for low and high metastatic efficacies were linked.

### Software and data availability

We are currently organizing our data and source code and will make both publically available as soon as possible (before journal publications). This data will include the raw image data, raw single cell images, and corresponding metadata, the trained autoencoder network, the feature representations of all cells, and code to perform our analyses.

### Funding and Acknowledgements

This work was supported by the Cancer Prevention and Research Institute of Texas (CPRIT R160622 to GD), the National Institutes of Health (R01GM071868 to GD; K25CA204526 to ESW), and the Israeli Council for Higher Education (CHE) via Data Science Research Center, Ben-Gurion University of the Negev, Israel (to AZ). We thank Sean Morrison for PDX-derived cell models. We thank Andrew R. Cohen for LEVER.

## Author Contribution

AZ, ESW and GD conceived the study. ESW designed the experimental assay, optimized culture conditions for PDX cells, and trained AN and JC to perform the experiments. AN performed all live imaging and mouse experiments. JC performed additional experiments. UE generated the PDX samples. AZ and ARJ developed analytic tools, analyzed, and interpreted the data. AZ drafted the manuscript. All authors wrote and edited the manuscript and approved its content. GD mentored all authors.

## Supplementary tables legends

**Table S1.** Panel of melanoma cell types used for this study: cell lines, melanocytes and patient-derived xenograft (PDX). Cells with high metastatic efficiency were derived from patients that exhibited metastases within 22 months, whereas cells with low metastatic efficiency were derived from patients that developed distant metastases within 22 to 50 months (Quintana et al., 2012).

**Table S2.** Genomics Variants in PDX and two cell lines. PDX-SNVs-Indel and CellLines-SNVs-Indels: Single nucleotide variants (SNVs) and Insertions/Deletions (Indel)s of high-quality variants identified were filtered for common variants in > 1% of any population in GnomAD. The information provided by the columns is as follows: Cell Type, corresponding sample labels in Table S1; Hugo Symbol, the HUGO Gene Nomenclature Committee approved gene name (symbol); Chromosome, the affected chromosome; Start, the mutation start coordinate; Variant_Classification translational effect of variant allele; Depth, the read depth across this locus in tumor BAM; RefCt, the Read depth supporting the reference allele in tumor BAM; AltCt, read depth supporting the variant allele in tumor BAM; Tumor_MAF, mutational allele frequency (AltCt/Depth) and GT, the genotype encoded as alleles values separated by either of “/” or “|”, e.g. The allele values are 0 for the reference allele (what is in the reference sequence), 1 for the ALT. PDX-CNVs and CellLines-CNVs: Copy number variants (CNVs) identified were filtered for genes mutated in > 5% of patients and > 10 patients in TCGA (Hoadley et al., 2018): The information provided by the columns is as follows: Cell Type, corresponding sample labels in Table S1; Hugo_Symbol, the HUGO Gene Nomenclature Committee approved gene name (symbol); Chromosome, the affected chromosome; Start, the mutation start coordinate; End, the mutational end position; Abberation is the type of copy number alteration as gain or loss; CN, the copy number of the gene; Score is calculated by CNV Kit as the sum of the weights of the bins supporting the CNV; Cytoband, the position on the chromosomal cytogentic band.

**Table S3.** Oncogenic Genomics Variants in PDX and two cell lines in commonly mutated genes (Hodis et al., 2012). The information provided by the columns is as follows: Annotation, the oncogenic effect, Hugo_Symbol, the HUGO Gene Nomenclature Committee approved gene name (symbol); Variant Type, CNV=Copy Number Variation, SNV=Single Nucleotide Variant and Indel= Insertion or Deletion; Variant, for SNVs and InDels, this is the HSGS protein change and is the aberration type (gain or loss) which has the predicted oncogenic effect; each following column is the Cell Type corresponding to Table S1 and the genotype or copy of the Cell Type.

**Table S4.** Non-oncogenic Genomics Variants in PDX and two cell lines in commonly mutated genes (ref landscape paper). The information provided by the columns is as follows: Hugo_Symbol, the HUGO Gene Nomenclature Committee approved gene name (symbol); Variant Type, CNV=Copy Number Variation, SNV=Single Nucleotide Variant and Indel= Insertion or Deletion; Variant, for SNVs and InDels, this is the HSGS protein change and is the aberration type (gain or loss) which has the predicted oncogenic effect; each following column is the Cell Type corresponding to Table S1 and the genotype or copy of the Cell Type.

## Supplementary video legends

**Video S1**. Heterogeneous morphology and local dynamics of patient derived melanoma cells. Shown are time lapse of single cells cropped from the field of view and used as the input for the autoencoder.

**Video S2**. Time lapse sequence of a representative field of view of m481 PDX cells.

**Video S3**. Reconstruction evolution and convergence during autoencoder training. Raw (top) and reconstructed (bottom) images during the autoencoder training minimizing the binary cross-entropy error between the two.

**Video S2Morphing**. Cell morphing in silico. Following the decoded cell image gradually morphing along an interpolated linear trajectory in the latent space between two cells.

**Video S5**. Morphing m498 PDX cell in silico from low to high metastatic efficiency by decoding the latent cell descriptor under gradual shifts in feature #56. The corresponding classifier’s score (“ClassifierScore”) and value of feature #56 in units of the z-score (“f56”) are shown.

**Video S6**. Morphing m405 PDX cell in silico from high to low metastatic efficiency by decoding the latent cell descriptor under gradual shifts in feature #56. The corresponding classifier’s score (“ClassifierScore”) and value of feature #56 in units of the z-score (“f56”) are shown.

**Video S7**. Morphing 100 m498 cells by gradually decreasing feature #56 (increasing classifier score).

**Video S8**. Time lapse of a m610 PDX cell spontaneously switching from the low to the high metastatic efficiency domain (as predicted by the classifier). Live imaging for 10 minutes.

## Supplementary figures

**Figure S1:**
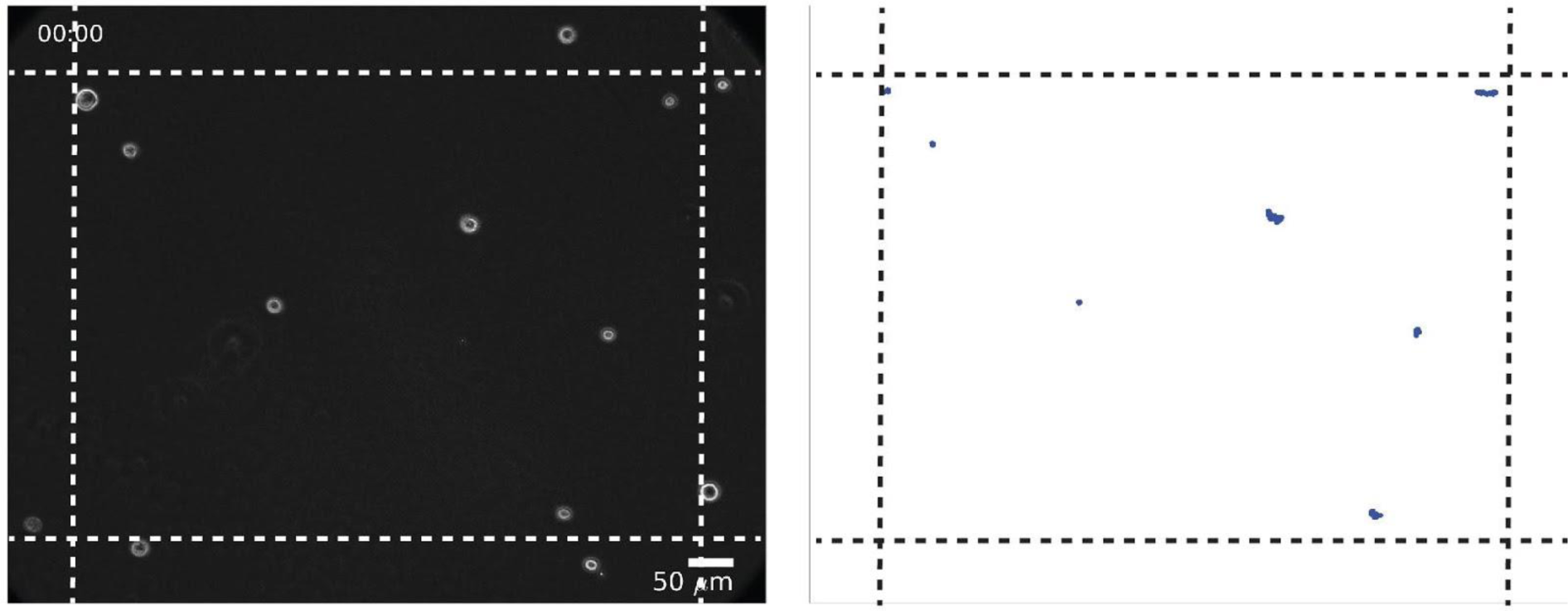
PDX melanoma on collagen were not migratory. Full field of view in phase contrast (left). Corresponding trajectories from 120 minutes indicate that cells are minimally motile (right). Only cells within the 70µm (dashed lines) were tracked (Methods).

**Figure S2:**
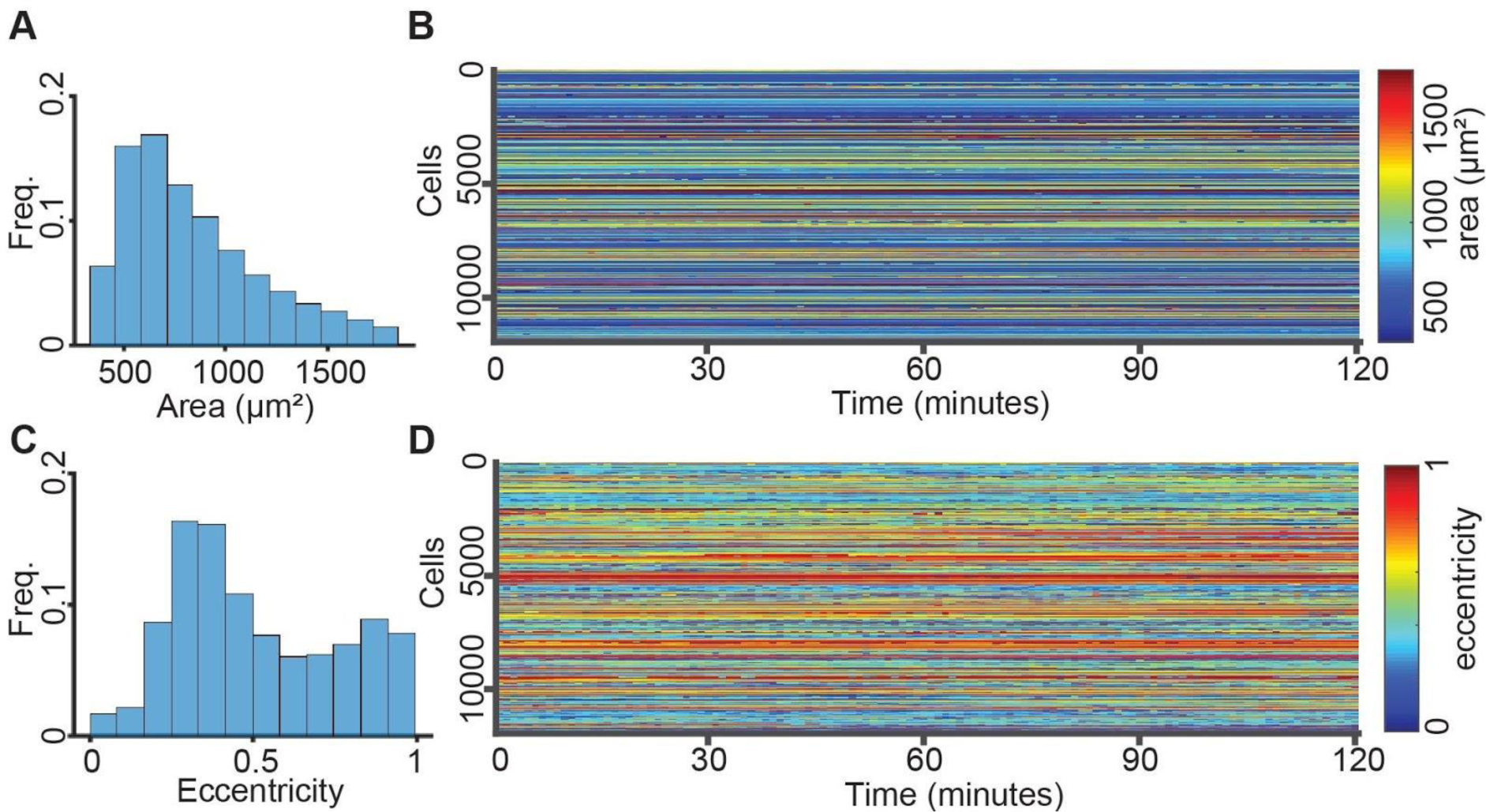
Quantitative visualization of cell shape over time. (**A-B**) Cell area. Distribution (**A**) and time-evolution (**B**) of all cells in all time points. (**C-D**) Cell eccentricity. Distribution (**C**) and time-evolution (**D**) of all cells in all time points.

**Figure S3:**
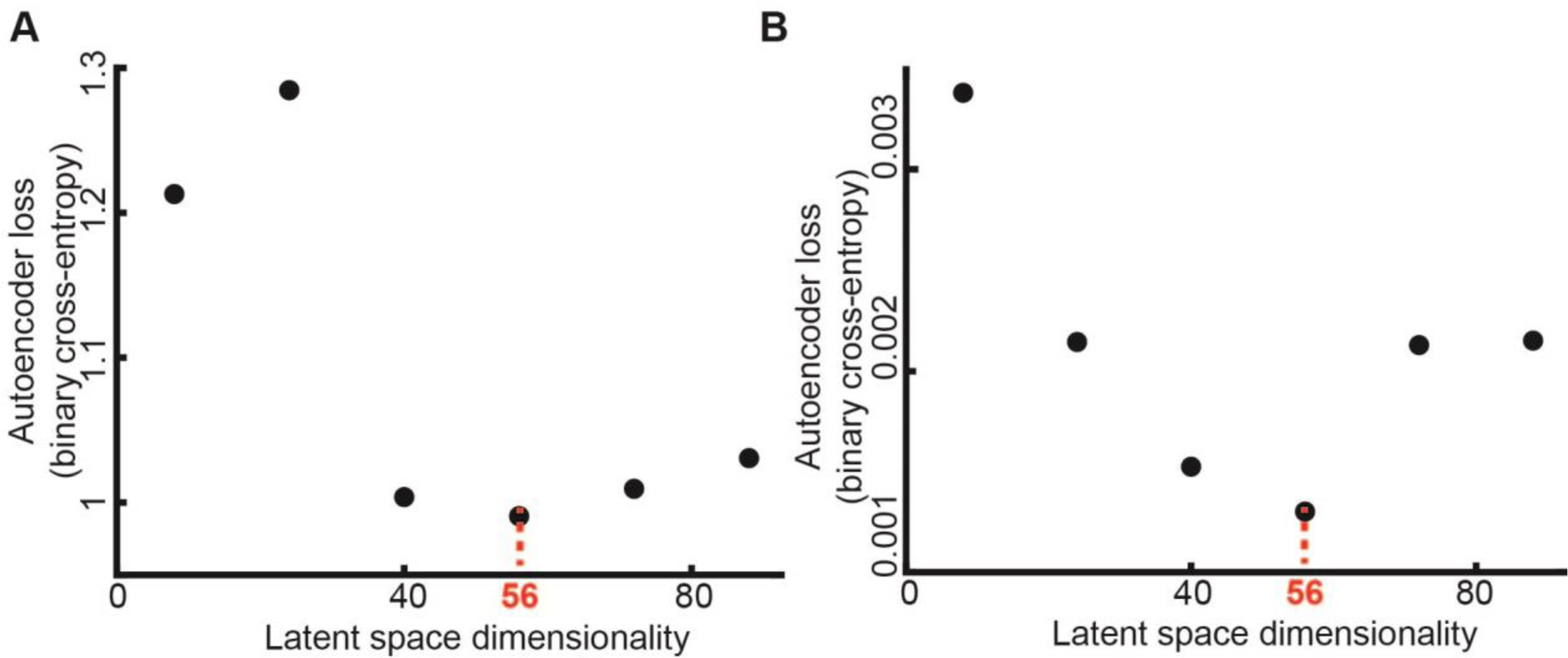
Loss and image reconstruction training error as a function of the latent space dimensionality. We selected the 56-dimensional latent vector based on minimizing loss and reconstruction error. (**A**) Autoencoder loss (binary cross-entropy) after training. (**B**) Mean square error for image reconstruction after training.

**Figure S4:**
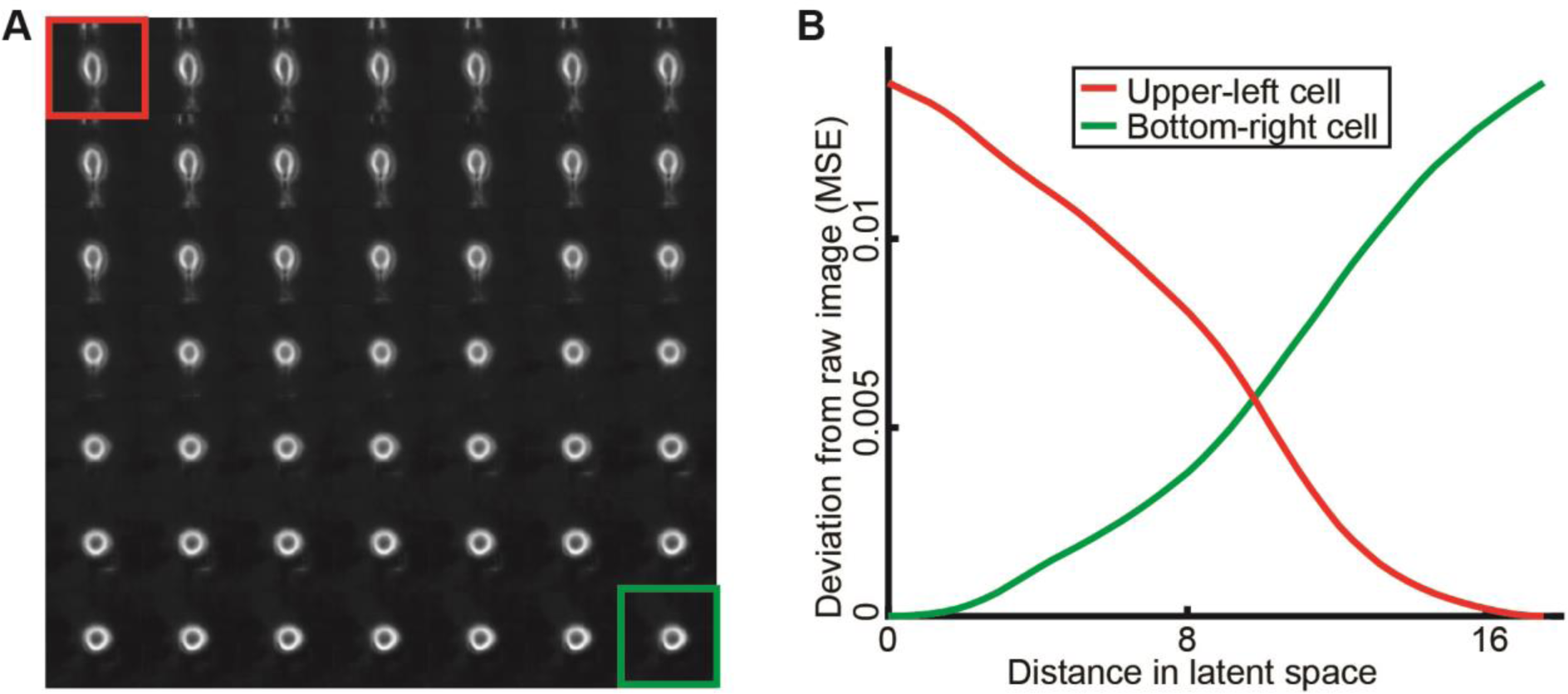
“Morphing” synthetic images between two random points in the latent space. (**A**) Series of cell image reconstructions along a straight line trajectory between two random points in the latent space (constant step size in latent space). The trajectory goes from top-left (red) to bottom-right (green). (**B**) Differences of images in panel A and to the start-(red) and endpoint (green) images.

**Figure S5:**
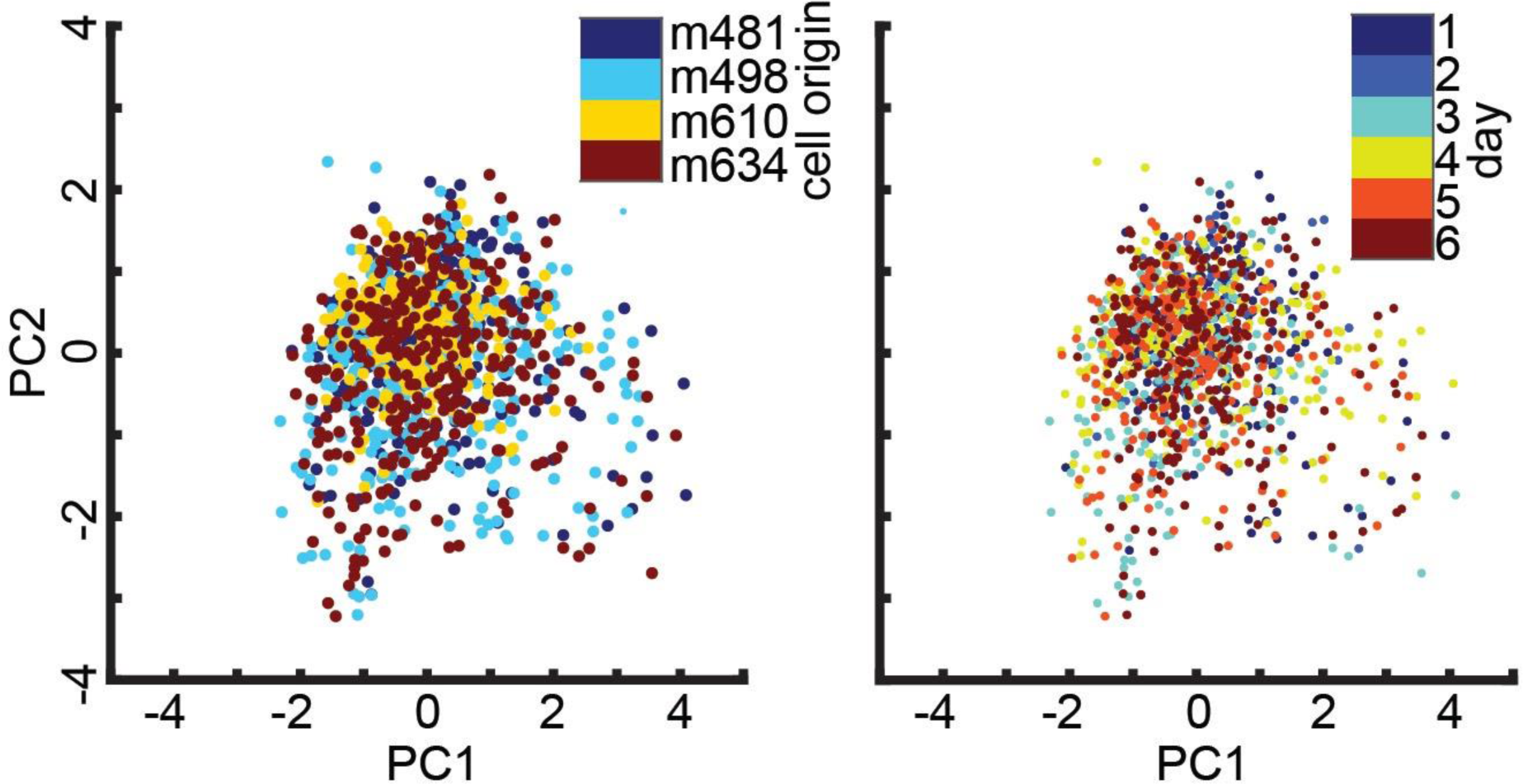
PCA projection of latent space descriptors of different PDXs on the same day (left) and of one PDX imaged on different days (right).

**Figure S6:**
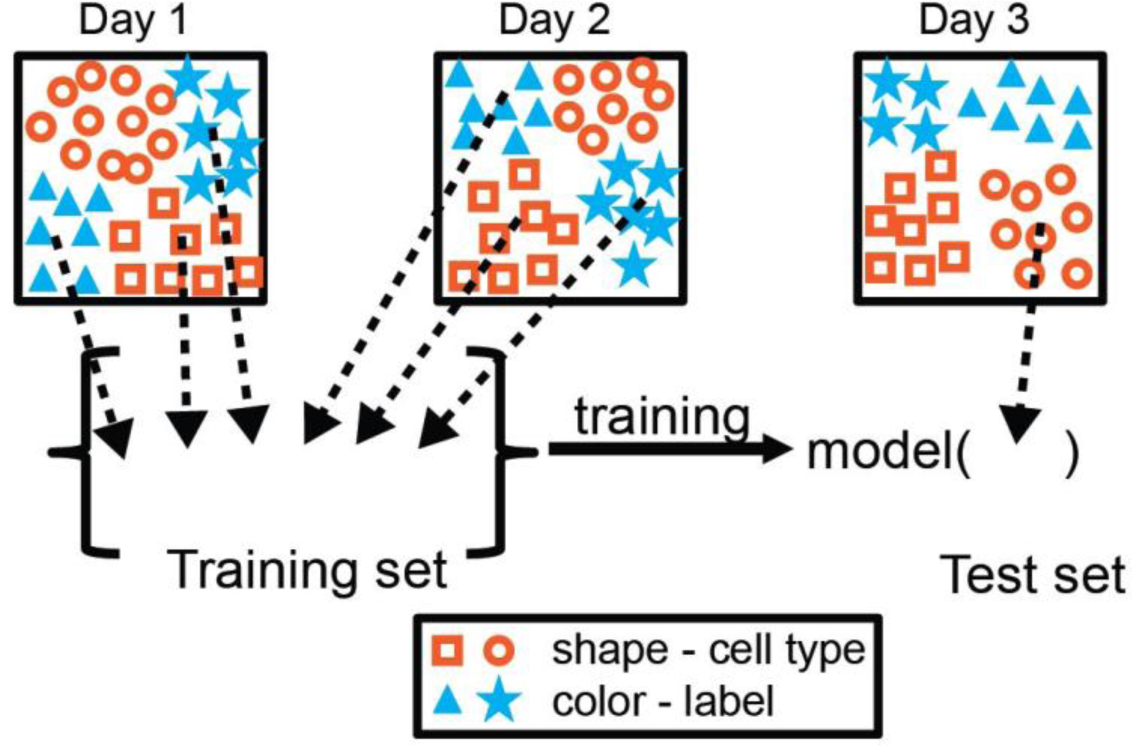
Blinding the cell type and the day of imaging. Multiple rounds of training and testing were performed. In each round, data from one cell type imaged in one day was used as the test dataset. The training set consisted of the remainder of the data, excluding the cell type at test and data from the same day of imaging. Thus, the trained model was completely blind to the test set. The model classified each cell in the test set, the overall mean classification accuracy for a specific cell type and imaging day was reported. The classifier’s score of every cell was recorded and accumulated for all cell type + imaging day pair for Receiver Operating Characteristic analysis. Besides excluding batch-effects by blinding the classifier to the day of imaging, this provided us with an increased number of observations (cell type, day) at the cost of a reduced number of cells per observation.

**Figure S7:**
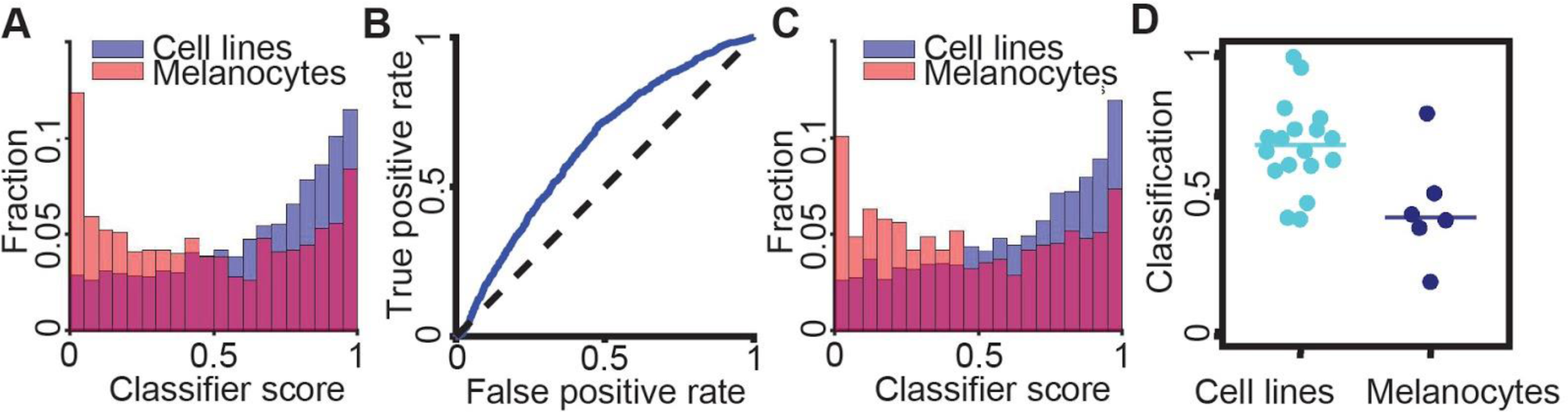
Discriminating melanoma cell lines from melanocyte lines. (**A**) Distribution of the single cell classifier score for classifiers blind to the <u>cell type</u> at test (Fig. 4A). (**B-D**) Discrimination results using classifiers that were blind to the <u>cell type and day of imaging</u> (Fig. S6). (**B**) Receiver Operating Characteristic (ROC) curve. AUC = 0.635. (**C**) Distribution of the single cell classifier score for classifiers blind to the cell type and day of the experiment at test (Fig. S6). (**D**) Accuracy in predicting the label ‘cell lines’ for a single cell as opposed to the label ‘melanocytes’. Each data point indicates the outcome (fraction of cells classified as ‘cell line’) of testing the cells of one melanoma cell line or melanocyte line on a particular day. N = 24: 18 cell lines, 6 melanocyte lines. 19/24 successfully predicted observations. Wilcoxon rank-sum test p = 0.026. Binomial statistical test p < 0.003.

**Figure S8:**
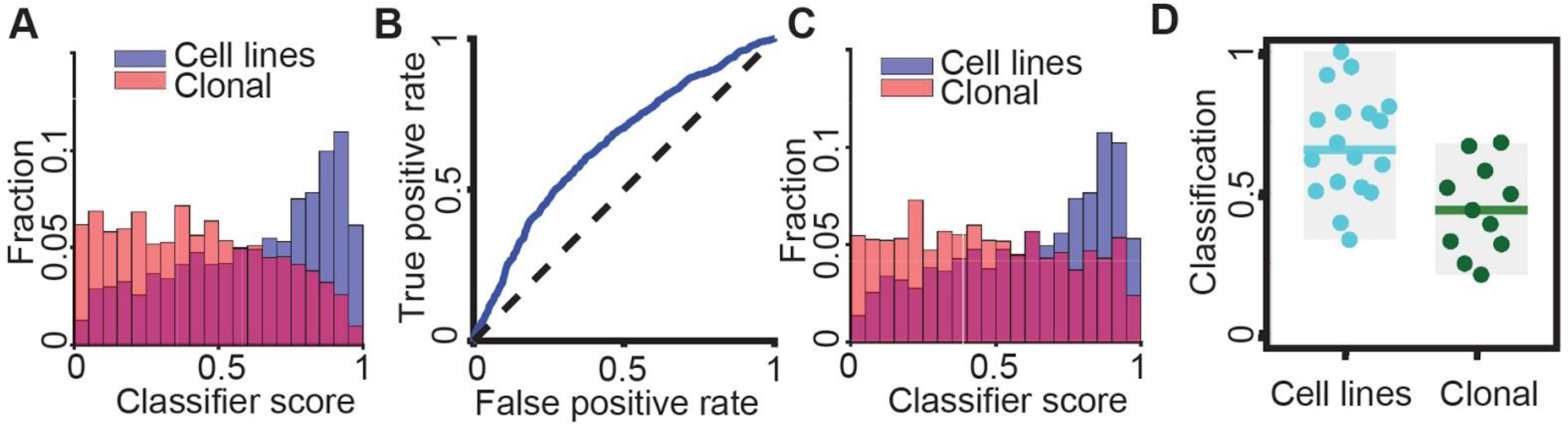
Discriminating melanoma cell lines from clonally expanded cell lines. (**A**) Distribution of the single cell classifiers score. Classifiers were blind to the <u>cell type</u> at test (Fig. 4A). (**B-D**) Discrimination results using classifiers that were blind to the <u>cell type and day of imaging</u> (Fig. S6). (**B**) Receiver Operating Characteristic (ROC) curve. AUC = 0.65. (**C**) Distribution of the single cell classifier score. Classifiers were blind to the cell type *and* day of the experiment at test (Fig. S6). (**D**) Accuracy in predicting the label ‘cell lines’ for a single cell as opposed to the label ‘clonal’. Each data point indicates the outcome of testing the cells of one melanoma cell line or clonal expansion line on a particular day. N = 29: 18 cell lines, 11 clonal expanded cells. 22/29 successfully predicted observations. Wilcoxon rank-sum test p = 0.0032. Binomial statistical test p < 0.0041.

**Figure S9:**
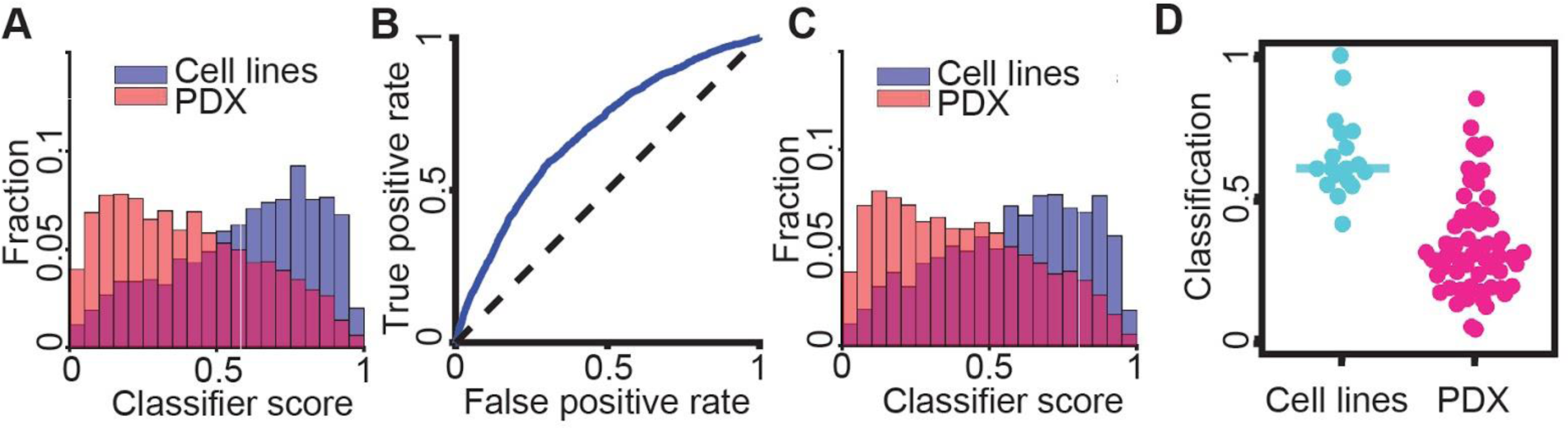
Discriminating melanoma cell lines versus PDXs. (**A**) Distribution of the single cell classifier score. Classifiers were blind to the <u>cell type</u> at test (Fig. 4A). (**B-D**) Discrimination results using classifiers that were blind to the <u>cell type and day of imaging</u> (Fig. S6). (**B**) Receiver Operating Characteristic (ROC) curve. AUC = 0.686. (**C**) Distribution of the single cell classifier score. Classifiers were blind to the cell type and day of the experiment at test (Fig. S6). (**D**) Accuracy in predicting the label ‘cell lines’ for a single cell as opposed to the label ‘PDXs’. Each data point indicates the outcome of testing the cells of one melanoma cell line or PDX on a particular day. N = 75: 18 cell lines, 75 PDXs. 63/75 successful predicted observations. Wilcoxon rank-sum test p < 0.0001. Binomial statistical test p < 0.0001.

**Figure S10:**
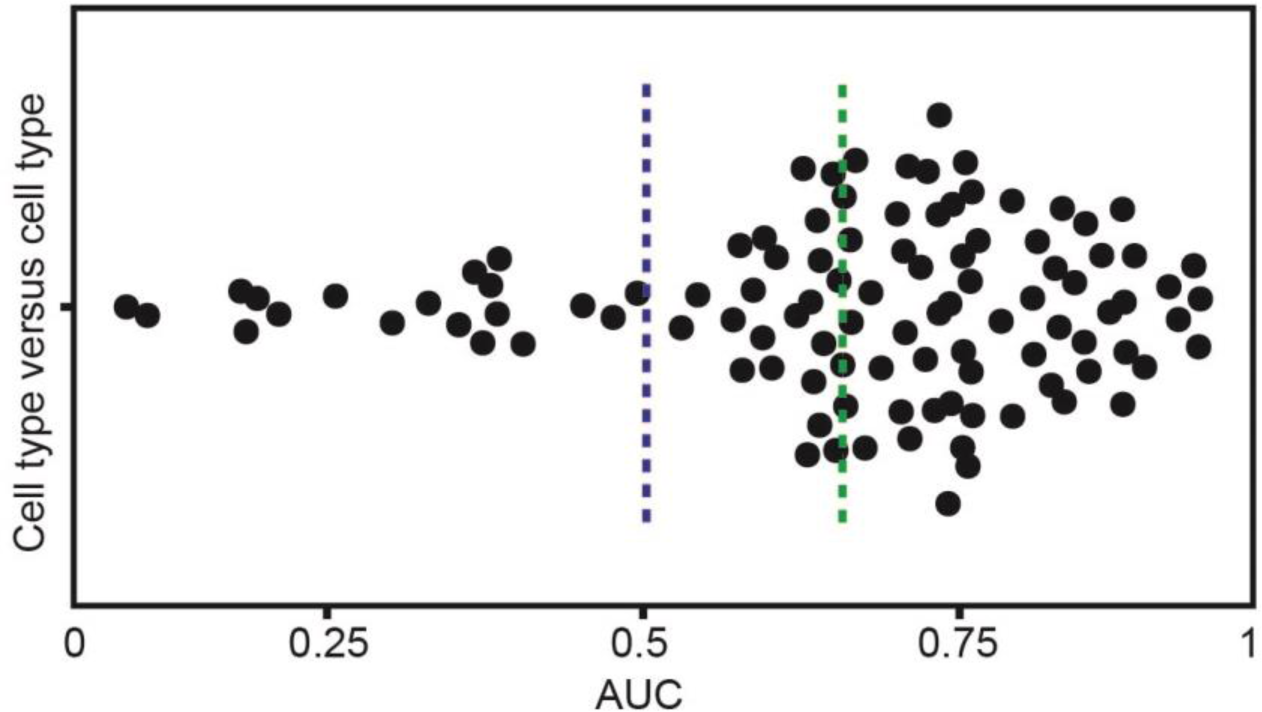
Pairwise discrimination of cell types. Discriminating two cell types from one another. Each data point indicates the AUC value for predicting the cell type label for single cells. Multiple rounds of training and testing were performed for each pairwise classification. In each round, data from one cell type imaged in one day was used as the test dataset, while the training set consisted of the remainder of the data, excluding data from the same day of imaging. Note that here the classifiers were blind to the day of imaging, but not to the cell type at test. The green dashed line is the mean AUC = 0.66. The blue dashed line indicates the AUC level of a random classifier. p-value < 0.0001 (Wilcoxon sign-rank test) rejecting the null hypothesis that pairs of different cell types cannot be discriminated.

**Figure S11:**
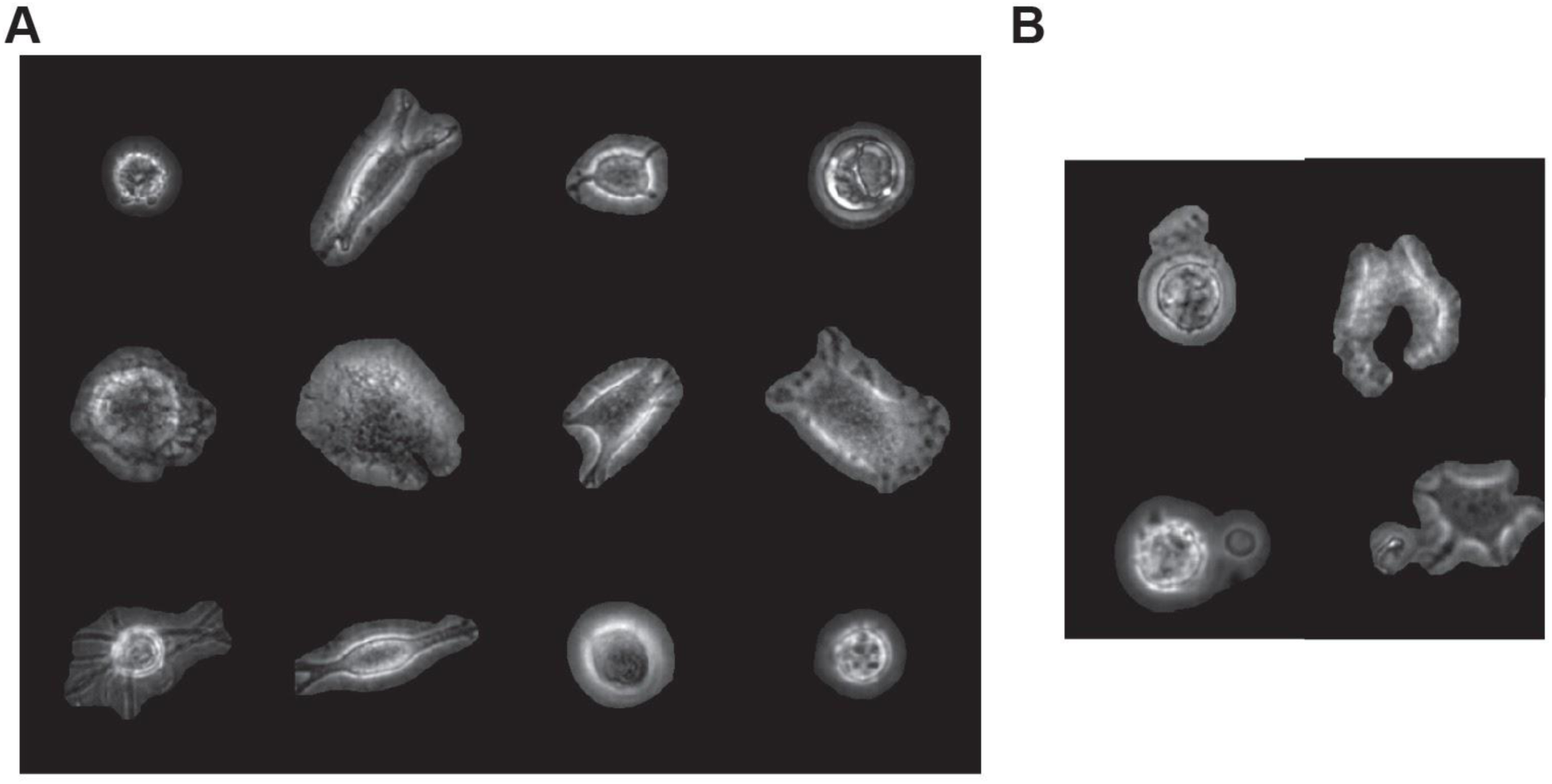
Single cell segmentation in phase-contrast images by LEVER. (**A**) Examples of successful segmentation. The region outside the segmentation mask is colored black. (**B**) Examples of failed segmentations.

**Figure S12:**
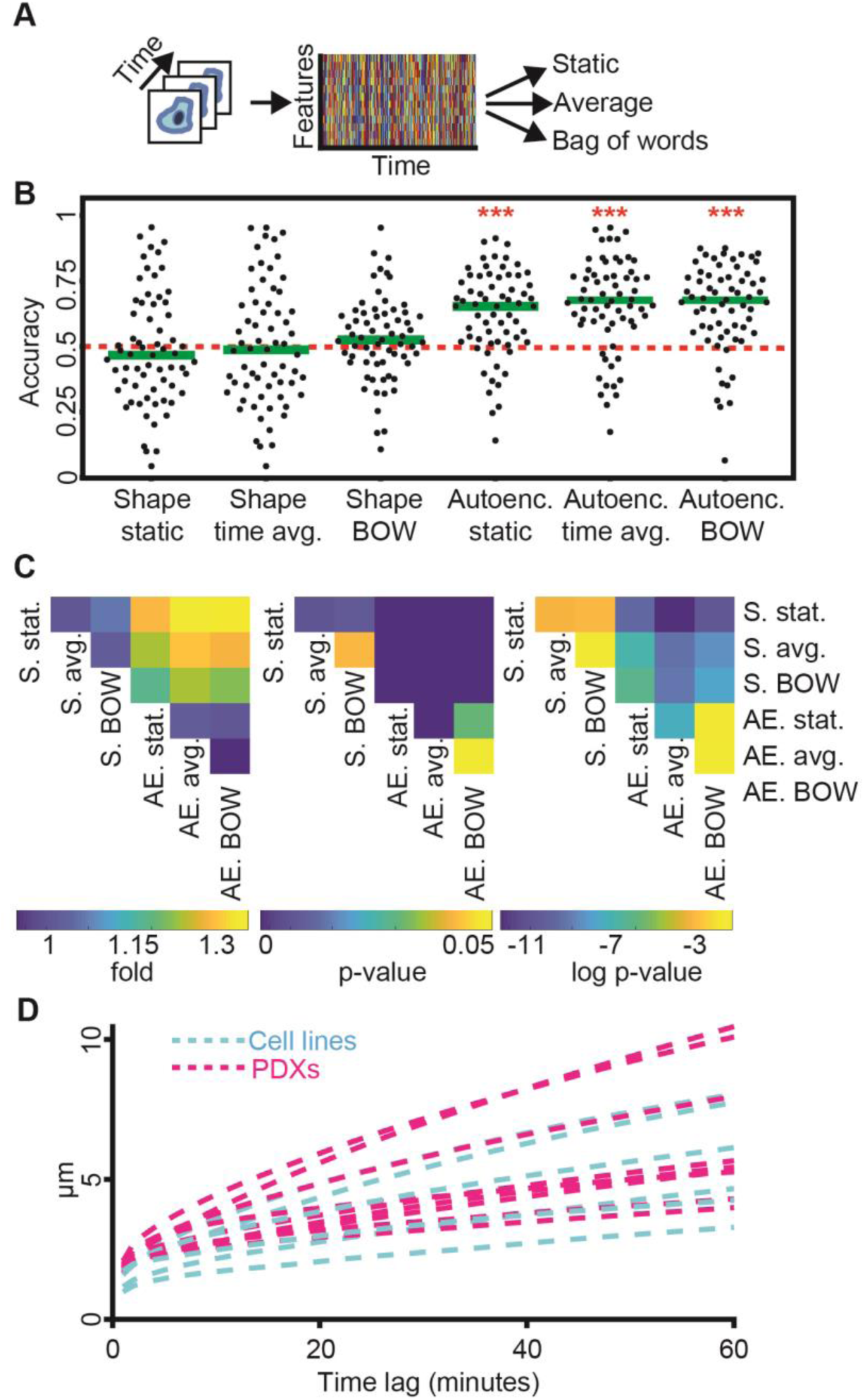
Classification comparison using cell shape and temporal information to distinguish cell lines from PDXs. (**A**) Three scenarios of incorporating temporal information in a cell descriptor applied to either cell shape-based features or latent space cell descriptors. (**B**) Accuracy in predicting the label ‘cell lines’ for a single cell as opposed to the label ‘PDXs’. Each data point indicates the outcome of testing the cells of one melanoma cell line or PDX on a particular day (Fig. S6). Classifiers derived from cell shape-based features could not discriminate between the two labels, regardless of the mode of incorporating temporal information. In contrast, the latent space cell descriptors slightly improved with explicit consideration of temporal information and all classifier modes significantly outperformed shape-based classifiers (*** - p-value < 0.0001, nonparametric Wilcoxon sign-rank test. N = 65 experiments of one cell type imaged in one day. The green line is the median. The dashed red horizontal line represents the random model.). (**C**) The latent cell descriptor outperforms shape features. Matrix visualization of the comparison of the different encodings. Fold (left), p-value (middle), log p-value (right, −3 corresponds to the p-value of 0.05). The average latent cell descriptor classification accuracy surpasses other cell encoding schemes. Stat - static, Avg. - average, BOW - bag of words. (**D**) Mean squared displacement analysis (MSD) analysis of single cell trajectories averaged over each cell type did not show discrimination between cell lines and PDXs. Maximal time lag of 60 frames (=minutes).

**Figure S13:**
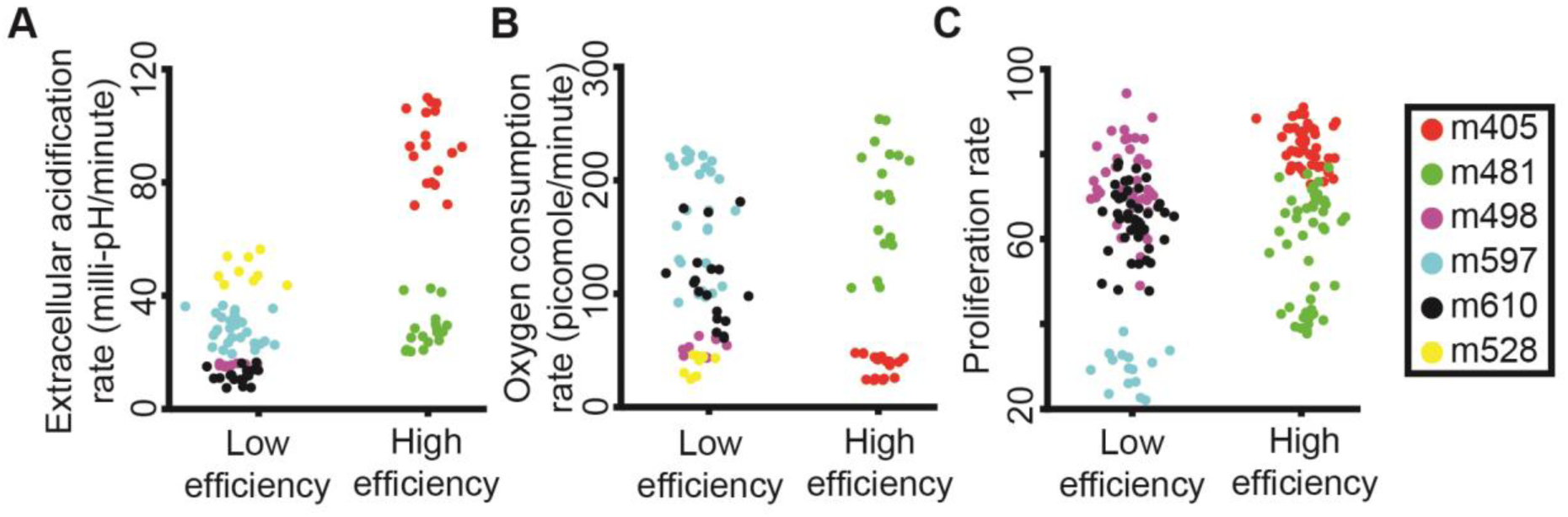
Standard assays of cell metabolic activity and proliferation cannot distinguish PDXs with low versus high metastatic efficiency. Each data point represents a technical replicate of the indicated assay. (**A**) Extracellular acidification rate. (**B**) Oxygen consumption rate. (**C**) Proliferation rate.

**Figure S14:**
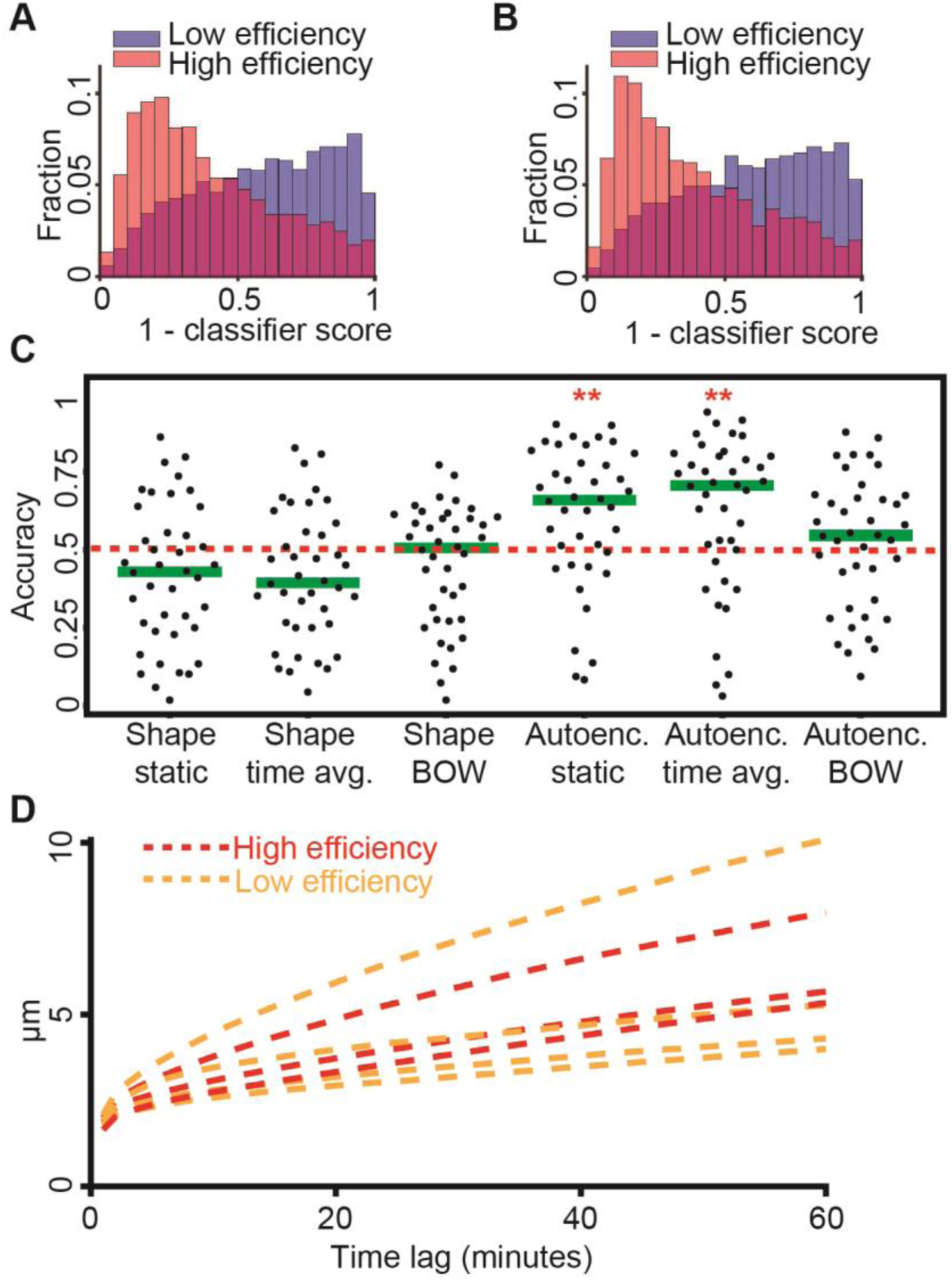
Discriminating high versus low metastatic efficient PDXs. (**A-B**) Distribution of the single cell classifier score. (**A**) Classifiers were blind to the cell type at test (Fig. 4A). (**B**) Classifiers were blind to the cell type and day of experiment at test (Fig. S6). (**C**) Accuracy of classifiers derived from shape based features and from latent space cell descriptors in predicting the label ‘low efficiency’ for a single cell. The classifiers include various modes of incorporating temporal information (Fig. S12). The 0.5 horizontal line reference the accuracy of a random classifier. Shape-based classifiers could not discriminate between PDXs with high and low metastatic efficiency. Classifiers derived from latent space cell descriptors performed significantly better than random ** - p-value < 0.01 (0.0053 for Autoenc. static, 0.0056 for Autoenc. time avg.), nonparametric Wilcoxon sign-rank test. N = 40 experiments of PDX imaged in one day. Green lines indicate medians of accuracy distributions. (**D**) Mean squared displacement (MSD) analysis of single trajectories averaged over each PDX could not distinguish between high and low metastatic efficiency. Max time lag of 60 frames (=minutes).

**Figure S15:**
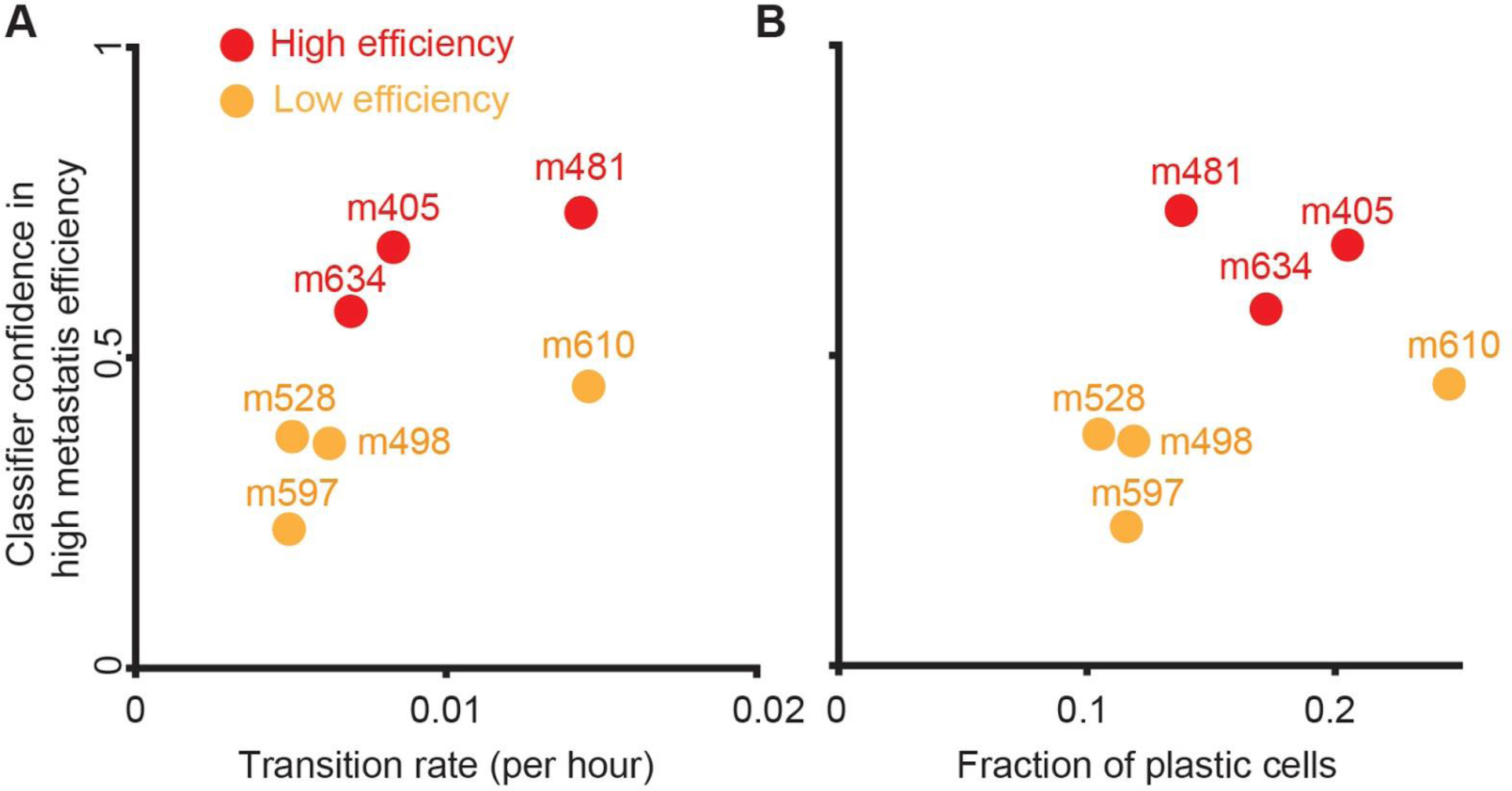
Predicted metastatic efficiency is correlated with cell plasticity. (**A**) Classifier score is correlated with PDX’s single cell’s transition rate from low-to-high or high-to-low. Pearson correlation coefficient of 0.58. (**B**) Classifier score is correlated with the fraction of a PDX’s single cells transitioning from low-to-high or high-to-low. Pearson correlation coefficient of 0.42.

**Figure S16:**
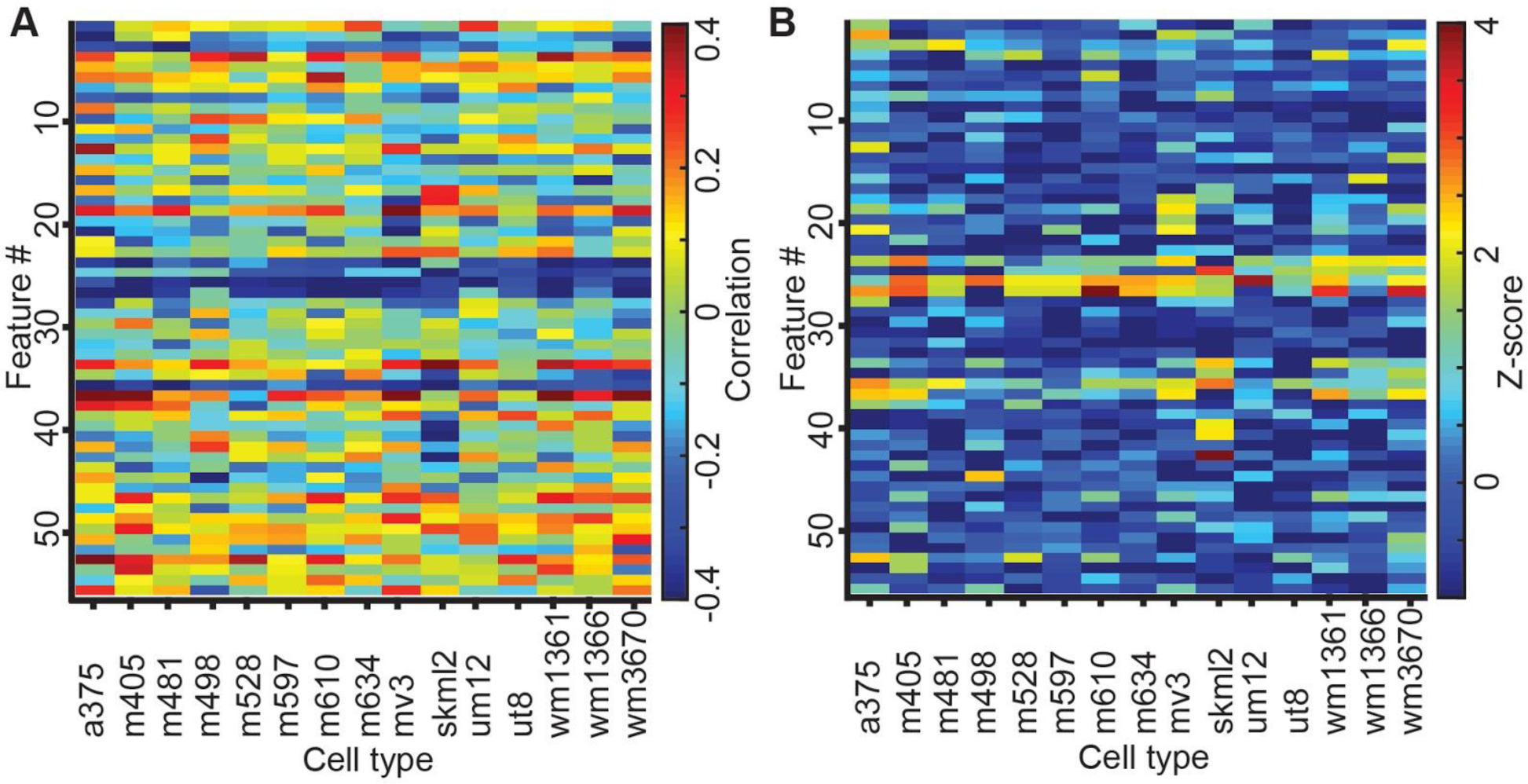
Multiple features are classification-driving for discriminating cell lines from PDXs. (**A**) Correlation values between all 56 features (y-axis) and the classifier scores for different cell types (x-axis). The correlation was calculated based on all cells from each cell type. (**B**) Normalized correlation values (Z-scores) between all 56 features (y-axis) and the classifier scores (x-axis) for different cell types.

**Figure S17:**
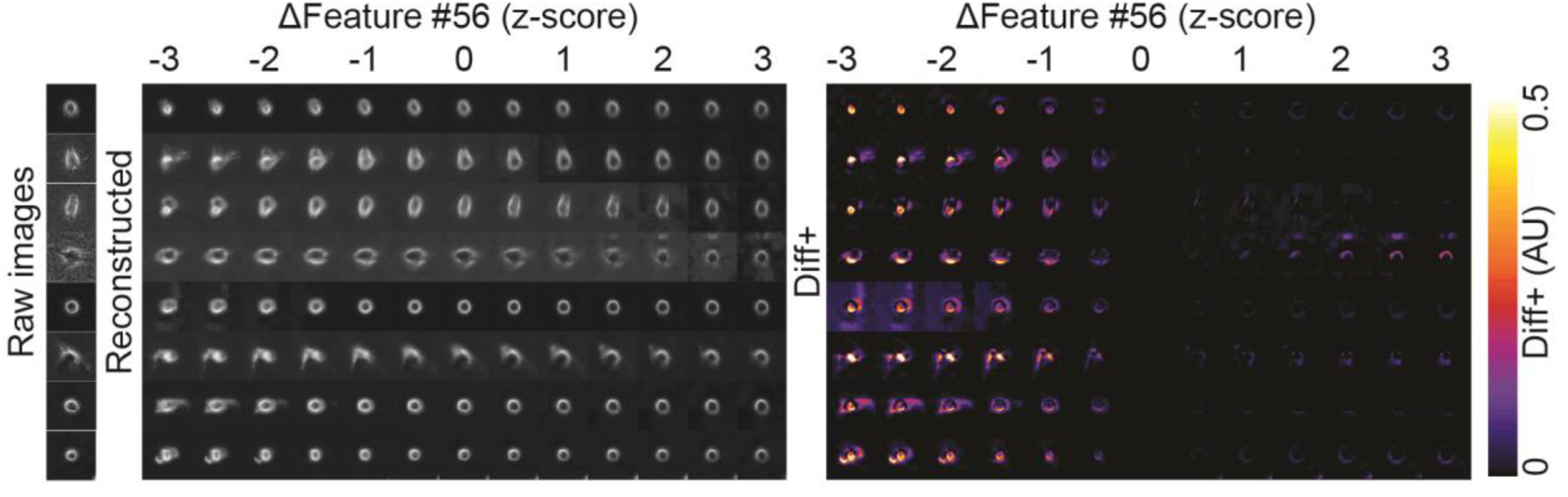
Panel of in silico cells generated by decoding a representative PDX cells’ latent space cell descriptor under gradual shifts in feature #56. Raw images (left), reconstructed images (middle), the positive values of the intensity differences between consecutive virtual cells (right).

**Figure S18:**
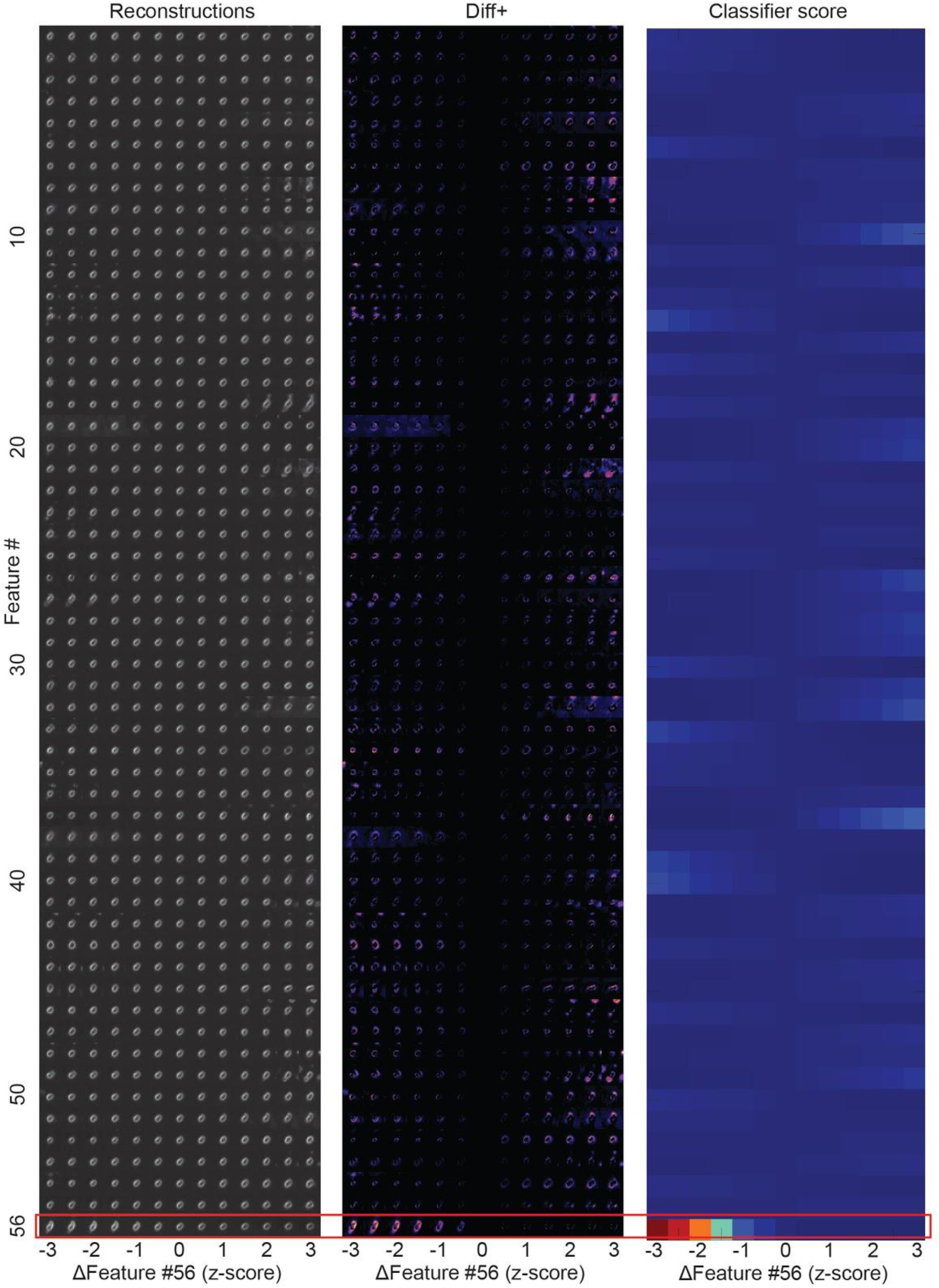
Visualization of in silico cells by altering each feature highlight the unique properties of feature #56. Reconstructed images (left), the positive values of the intensity differences between cells with different values in feature #56 (middle), the classifiers’ predicted scores (right).

**Figure S19:**
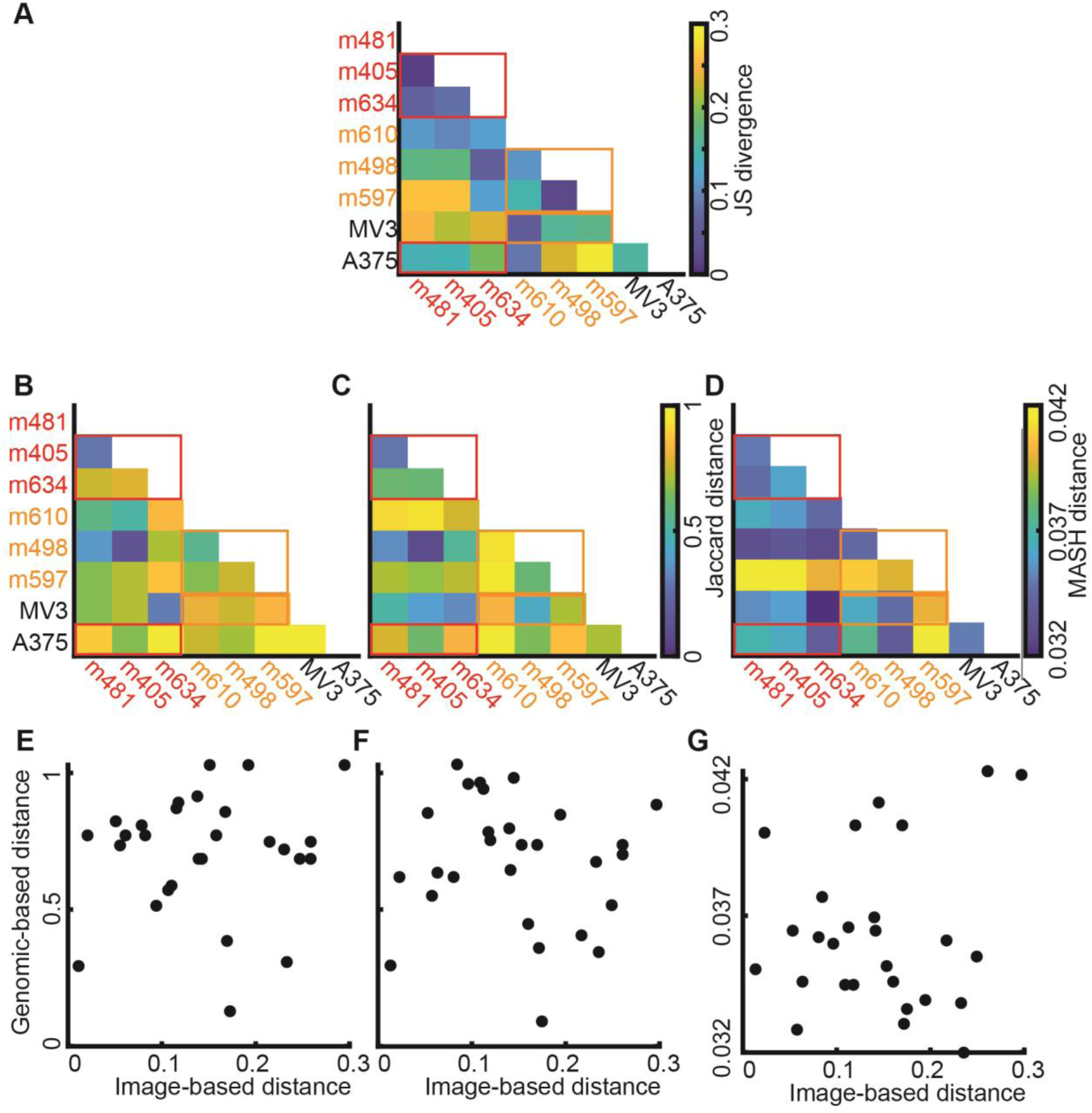
Genomic markers could not distinguish between high and low metastatic efficiency. Distance measures between pairs of different cell types were compared between image-based (classifier scores) and genomic-mutational based information. (**A-D**) Distance matrices between pairs of cell-types. Red sub-matrices indicate the distances between PDXs (and the A375 cell line) classified as highly metastatic. Orange sub-matrices indicate the distances between PDXs (and the MV3 cell line) classified as highly metastatic. (**A**) Distance matric derived from image based classifier scores. Individual distances were computed based on the Jensen-Shannon divergence of the classifier score distributions for single cells in each of the compared cell types. The sub-matrices of cell types with similar levels of metastatic efficiency show low distances compared to matrix bins comparing cell types with differing metastatic efficiency. (**B-D**) Distance matrices derived from the genomic profiles of cell types cannot distinguish between high and low metastatic efficiency. (**B**) Distances calculated based on the Jaccard index of the mutational state of the oncogenic mutations in the 20 top mutated genes in melanoma. (**C**) Distances calculated based on the Jaccard index of the non-oncogenic mutations from those same genes. (**D**) Distances calculated by application of the alignment-free method MASH to the sequences from the entire 1400 gene panel. (**E-G**) Distances derived from image-based versus genomics-based cell-type to cell-type distinction are not correlated. Each datum holds the matched pair classifier- and genomic-distances between two cell types. E, F, and G correspond to the matrices in B, C and D, each correlating with the distance matrix in A. No correlation was found to be statistically significant.

**Figure S20:**
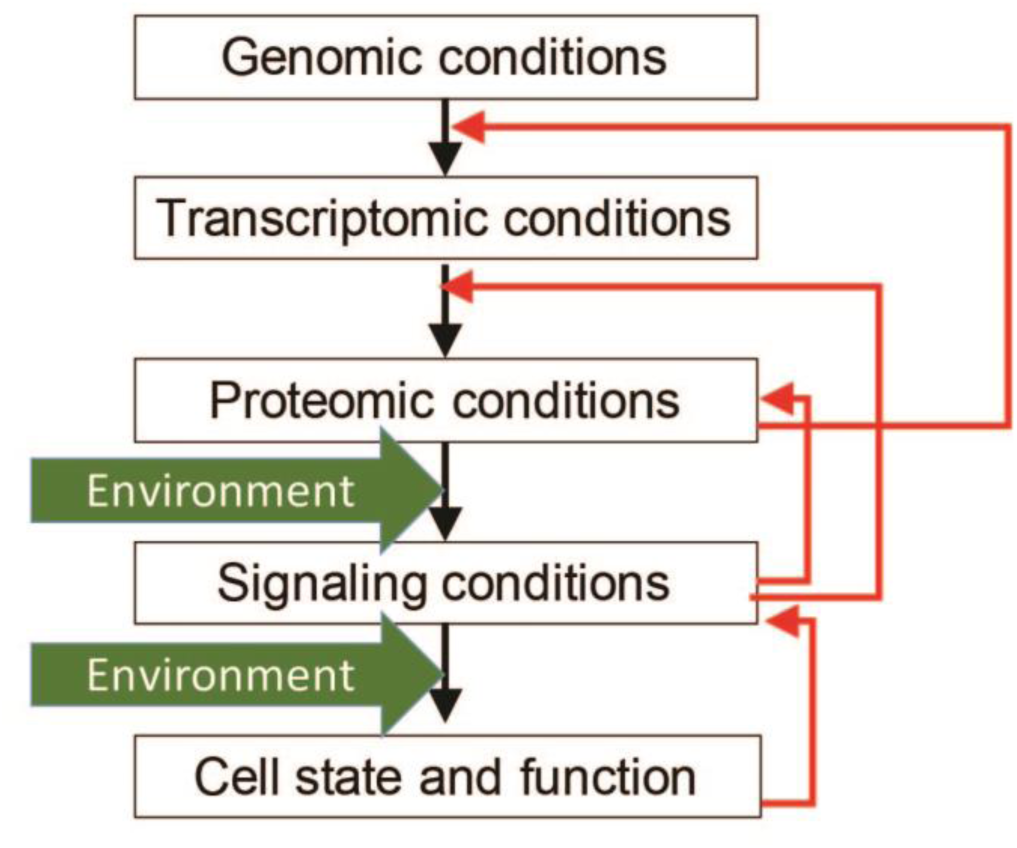
Hierarchical link between the genomic condition and cell function. In many scenarios of cancer the linear link (block) is intercepted by feedbacks (red). For example, cell morphology is known to affect signaling; signaling affects the proteomic condition directly and indirectly through post-translational modifications and regulation of proteostasis; and the proteomic conditions affect gene expression via transcriptional regulation. Dependent on the balance between forward and feedback interactions a particular genomic condition does or does not causally relate to cell functions like metastasis. Accordingly, genomic readouts may or may not be adequate predictors of the disease outcome. The relationship between genes and cell function is further weakened by influences of the environment.

